# Genetic basis of expression and splicing underlying spike architecture in wheat (*Triticum aestivum* L.)

**DOI:** 10.1101/2023.05.04.539218

**Authors:** Guang Yang, Yan Pan, Licao Cui, Mingxun Chen, Qingdong Zeng, Wenqiu Pan, Zhe Liang, Dave Edwards, Jacqueline Batley, Dejun Han, Pingcheng Deng, Hao Yu, Robert J Henry, Weining Song, Xiaojun Nie

## Abstract

**Introduction:** Wheat is one of the most important staple crops worldwide, and an important source of human protein and mineral element intake. Continuously increasing stable production of wheat is critical for global food security under the challenge of population growth and limited resource input.

**Objective:** Spike architecture determines the potential grain yield of wheat. However, the mechanisms of transcriptional regulation of spike architecture in wheat remain largely unknown, limiting further genetic improvement of wheat yield. In this study we explored the genetic basis of spike architecture in wheat.

**Methods:** Population RNA-seq methods were used to identify the eQTLs and sQTLs associated with spike architecture and applied this to dissection of the genetic basis of gene expression and splicing controlling these complex yield-related traits.

**Results:** In total, 4,143 expression quantitative trait loci (eQTLs) and 12,933 splice QTLs (sQTLs) were identified in wheat based on 178 RNA-seq samples, revealing 774 cis-eQTLs and 321 cis-sQTLs for 86 eGenes and 73 sGenes, respectively. Integration of eQTLs and sQTLs with genome-wide association study (GWAS) identified dozens of additional novel candidate genes that may contribute to spike-related traits. Gene network analysis showed that eQTLs and sQTLs were widely involved in the co-expression modules that regulate wheat spike architecture. Notably, the eQTL locus *AX-108754757* regulated the expression of 5 eGenes that negatively controled grain number per spike. *AX-111592099* regulated both the splicing and expression of *TraesCS7B02G442100,* encoding an E3 ubiquitin ligase, and playing a central role in regulating spike length.

**Conclusion:** This study provides new insights into the genetic basis of spike architecture. This improved understanding of spike-related traits in wheat will contribute to more rapid genetic improvement.

## Introduction

Wheat is one of the most important staple crops worldwide, accounting for approximately 30% of the global cultivated area, and providing 20% of the world’s food consumption [1]. Wheat is also an important source of human protein and mineral element intake [2]. Stable but increasing production of wheat is critical for global food security in response to growing demand due to population increases and despite limits to resources for agricultural production. It is estimated that wheat production needs to be increased by 1.5% annually to meet the demands of 9 billion people in 2050 [3]. Consequently, improvement of wheat yields is urgent.

Similar to other cereal crops, such as maize, rice and barley, the yield of wheat per plant is a complex trait determined by spike number per plant, grain number per spike (GN) and one-thousand kernel weight (TKW), all of which depend largely on spike architecture and are thus considered as spike-related traits. For example, the spikelet number (SN) and GN are influenced significantly by spike length, and TKW is mainly regulated by grain length (GL), grain width (GW) and grain length-to-width ratio [4]. Moreover, GN and TKW are the major targets for improving wheat yield [2].

Therefore, the identification of genetic loci and causal genes controlling these spike-related traits not only holds the promise of elucidating the genetic basis of wheat spike architecture, but also lays a foundation for the improvement of wheat yield. As complicated quantitative traits, wheat spike-related traits are controlled by multiple genes, and also affected by non-genetic confounding variables [5]. In light of their importance, extensive studies have been performed to identify the genes regulating spike architecture and spike-related traits, and some breakthroughs have been achieved. The *Q* locus, encoding an AP2-like transcription factor, is one of the best known and widely studied domestication genes regulating spike compactness, brittle rachises and grain morphology [6]. Feng *et al* found that mutation of *ARGONAUTE1d* resulted in shorter spikes and fewer grains per spike in durum wheat [7]. At the same time, GWAS and QTL analysis have also been widely applied in detecting genetic and phenotypic associations to dissect similar complicated quantitative traits in wheat, reducing the influence of confounding factors on complex traits by using a mixed linear model [8,9]. QTL mapping based on an F2 population of ZM366 and LH6 was performed to identify 48 QTLs for six spike-related traits [10]. A total of 120 loci were shown to be associated with spike-and yield-related traits using SNP-GWAS and haplotype-GWAS [11]. However, the genetic mechanisms, especially the molecular modules, associated with the development of the wheat spike, the complexity of this genetic regulation, and association with grain yield is still not well understood. Identification of the causal genesthat affect these phenotypic traits using the GWAS approach remains highly challenging [12,13].

Analysis of gene expression is an important way to understand biological functions, and identify the genetic variants that influence complex traits variants by affecting gene expression. eQTLs can be mapped and leveraged to link genome variants with target gene expression levels and have begun to be used to mine causal genes [14]. Walker et al (2019) identified and analyzed gene expression variation in the human brain, so as to discover causal genes closely related to autism spectrum disorder (ASD) and schizophrenia (SCZ), which are difficult to identify in common association analysis [15]. Pavlides et al (2016) combined GWAS and eQTL to obtain 28 important genes controlling complex traits in the human body, including some newly identified genes that could not be found through general association analysis [16]. In maize, 25,660 eQTLs for 17,311 genes were identified through RNA-seq of the RIL population, and bHLH transcription factor R1 and hexokinase HEX9 were demonstrated to function as crucial regulators of flavonoid biosynthesis and glycolysis [17]. Furthermore, 97 genes that were associated with drought tolerance, of which 6 major haplotypes of *abh2* were verified by the CRISPR-Cas9 approach in maize, were detected by eQTL analysis [18]. In addition, alternative splicing (AS) is a vital regulatory mechanism that enriches genetic control and affects gene function at the post-transcriptional level, and is also regulated by genetic variations (sQTLs) [19]. Chen *et al* (2018) reported the genome-wide identification of AS variants in maize and the effects of AS variation on gene function and phenotype variations, and found that *ZmGRF8* was influenced by a genetic factor (cis-sQTL), resulting in the transformation of transcriptional isoforms and a change in miRNA binding sites [20].

Analysis of gene expression and alternative splicing regulation provide an effective approach to bridge the gap between genetic variations and phenotypic traits, not only to enable the efficient mining of the causal genes but also to help identify the gene regulatory networks underlying complicated traits. However, few studies on gene expression and splicing variations of complex traits in wheat, especially spike-related traits have been reported to date.

To reveal the genetic basis and molecular modules of wheat spike architecture, a total of 178 RNA-seq data sets from the developing spike collected from 89 elite wheat varieties in two environments were generated in this study. Based on 660K SNP genotyping, eQTL and sQTL loci were first identified for the wheat spike. Then, a co-expression network was constructed, and the modules associated with 12 spike-related traits were identified. By integration of eQTLs and sQTLs with GWAS signals, we obtained the causal genes and molecular modules underlying spike-related traits (Figure 1). This study demonstrates that the rapid identification of the genetic basis of complicated agronomic traits is feasible through eQTL and sQTL analysis and also contributes to molecular breeding for yield improvement in wheat and beyond.

**Figure 1.**
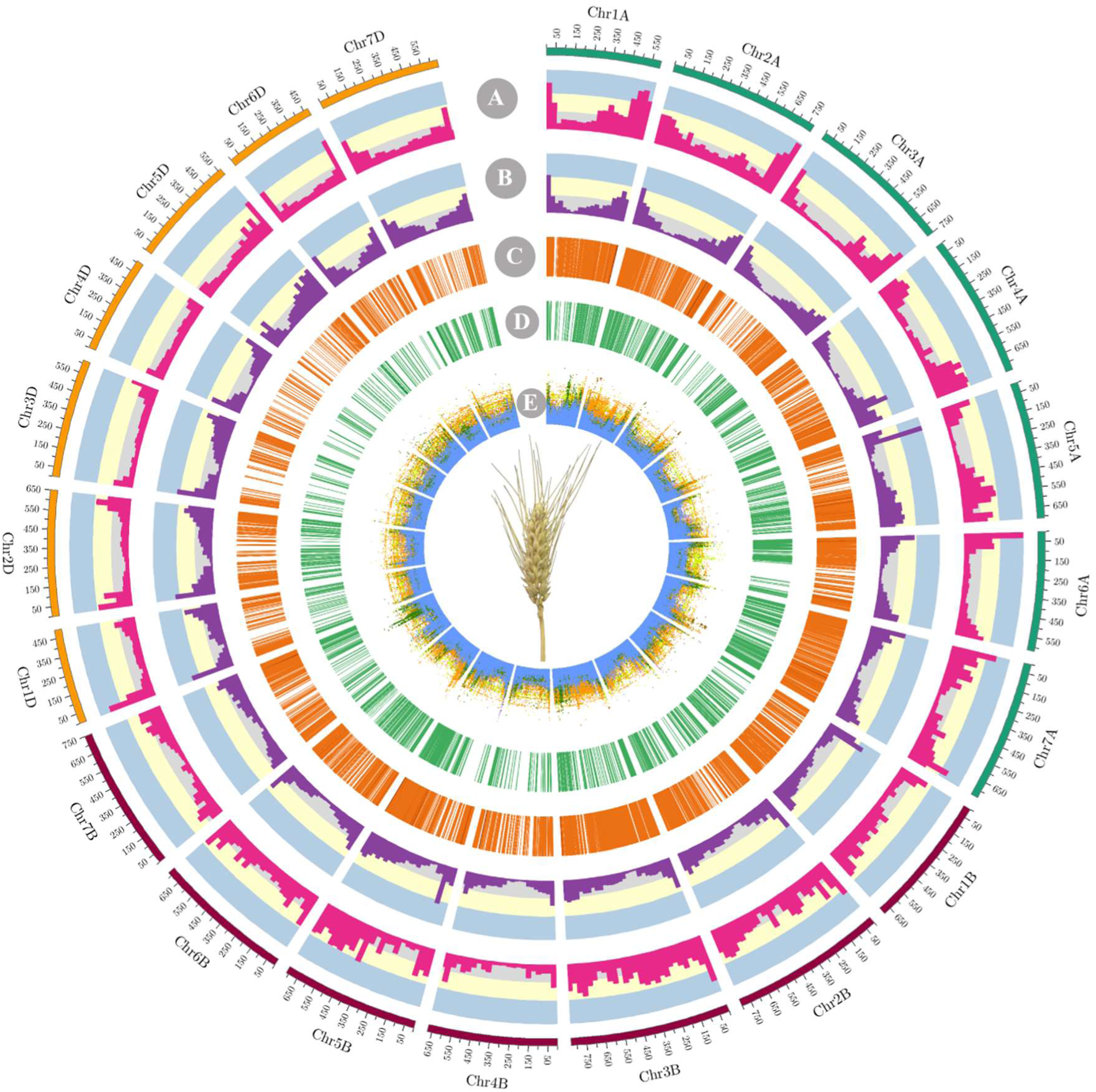
Summary of GWAS, eQTL and sQTL analysis of spike-related traits in wheat. A, gene density; B, SNP density; C, sQTLs; D, eQTLs; E, GWAS signals of spike length (SL), grain number per plant (GN), spike number per plant (SN), grain roundness (GR), grain diameter (GD), grain circumference (GC).

## Results

### Generation and characterization of a multi-omics dataset

#### Transcriptome assembly and variation among varieties

RNA extracted from spike samples at the heading stage was collected from a total of 89 wheat varieties grown under two environmental conditions, and subjected to RNA-seq. We obtained high quality reads (Figure S1A), which were further mapped onto the IWGSC_V1.1 reference genome, and 33.01 million read pairs per sample were uniquely mapped to the genome, with an average 95.2% mapping ratio and 96.0% genome coding region coverage (Figure S1B and 1C). Totally, 117,624 (novel: 6,834; known: 110,790) genes with 287,457 (novel: 150,401; known: 137,056) transcripts were detected, of which 50,493 genes possessed alternative splicing isoforms.

Compared with the reference genome, the splicing ratio of the novel assembled transcripts increased by 27.3% (Figure S1D), suggesting a more comprehensive transcriptome landscape was obtained.

#### GWAS analysis of spike (grain)-related traits

Genotype analysis of the above varieties was performed using the wheat 660K assay, and a total of 590,136 quality-filtered SNP loci (SNPs) were obtained and employed to perform GWAS analysis for 12 spike-related traits, including spikelet number (SN), grain number (GN), spike length (SL), grain area (GA), grain roundness (GR), grain circumference (GC), grain diameter (GD), ratio of grain length to grain circumference (GL/GC), ratio of grain width to circumference (GW/GC), ratio of grain length to width (GL/GW), ratio grain area to grain circumference (GA/GC), and thousand kernel weight (TKW). A total of 3,231 over-represented signals were obtained, with 263 signals associated with SN, 611 with GN and 859 with SL (Table S1), of which 652 (23.4%) out of 2,789 unique SNPs were located within the annotated gene regions. Among them, *TraesCS6A02G194600* was associated with GN, which is orthologous to *OsFUWA* that encodes an NHL domain-containing protein and alters panicle architecture, grain shape and grain weight by restricting excessive cell division (Table S2; Figure S2) [21]. *TraesCS7A02G393800* was associated with SL, and is orthologous to *AtCSLA9* involved in seed germination and fruit development [22]. Overall the GWAS results provided an indispensable resource for downstream analysis and mining of causal genes.

#### Gene co-expression analysis

Using the criteria described in the Method section, we further constructed co-expression networks based on the filtered genes to obtain 25 co-expression molecular modules (Figure S3A). The paired module comparison revealed that modules M16 and M4, M12 and M19, M22 and M15 showed a strong correlation with each other (Figure S3B). Based on module-trait associated analysis, the potential relationships between the gene modules and traits were further characterized (Figure S3C). To further uncover the biological functions of these modules, we performed Gene Ontology (GO), Plant Ontology (PO) and Trait Ontology (TO) analysis (Table S3) and found that many grain and spike related terms were enriched by these modules. For example, ubiquitin catabolic process (GO:0006511), with potential roles in grain and spike development, was enriched in the M1, M2, M9, M12 and M14 modules. Protein ubiquitination (GO:0016567) was enriched in the M5, M9 and M13 modules. The glume term (PO:0009039) was enriched in M3 and M1 modules; plant embryo axis term (PO:0019018) was enriched in the M4 module; and plant ovary related terms were enriched in the M4 (PO:0005022), M1 (PO:0009072) and M6 (PO:0025272) modules. These molecular modules, with their potential functions, suggested that the gene co-expression was related to wheat spike-related traits.

#### Identification of sQTLs and eQTLs

A gene splicing genome-wide association study (sGWAS) and gene expression genome-wide association study (eGWAS) were further performed based on the gene alternative splicing ratio and expression level. Local sQTLs and eQTLs were defined as the linked genes that were situated in a supported region with a LD value r^2^ > 0.2 [20, 23], with others considered as distant QTLs. Thus, the threshold for local sQTL-and eQTL-linked intervals was set to 11.396 MB (Figure S4). To conduct the sGWAS, alternative splicing categories were first predicted. Consistent with a previous study in maize (Chen *et al.*, 2018), intron retention ranked as the most abundant type, followed by alternative acceptor sites (AA: 20.0%), alternative donor sites (AD: 12.5%) and exon skipping (ES: 7.3%) (Figure 2A). We found that 12,933 sQTLs, including 170 (1.3%) cis-sQTLs, 12,612 (97.5%) trans-sQTLs and 151 (1.2%) cis/trans-sQTLs, were associated with the splicing ratio of 690 genes and 857 transcripts (threshold: P < 6.83 × 10^−8^) (Figure 2B; Table S4). Remarkably, the median splicing ratio was 8.0% and 2.5% for cis-and trans-sQTLs, respectively (Figure 2C; Figure S5A, B), suggesting cis-sQTL may play the key role in splice pattern domination and transcriptional regulation during wheat spike development. eGWAS analysis identified 4,143 eQTLs that were associated with the expression level of 286 genes (threshold: P < 6.83 × 10^−8^) (Figure 2D; Table S5). Among them, 3,369 (81.32%) eQTLs were trans-eQTLs, 731 (17.64%) were cis-eQTLs and 43 (1.04%) were both trans-eQTLs and cis-eQTLs. Among the 286 eGWAS-associated genes, the expression level of 19 genes was associated with cis-eQTLs, that of 200 genes was associated with trans-eQTLs, and the remaining 67 showed associations with both cis-and trans-eQTLs.

**Figure 2.**
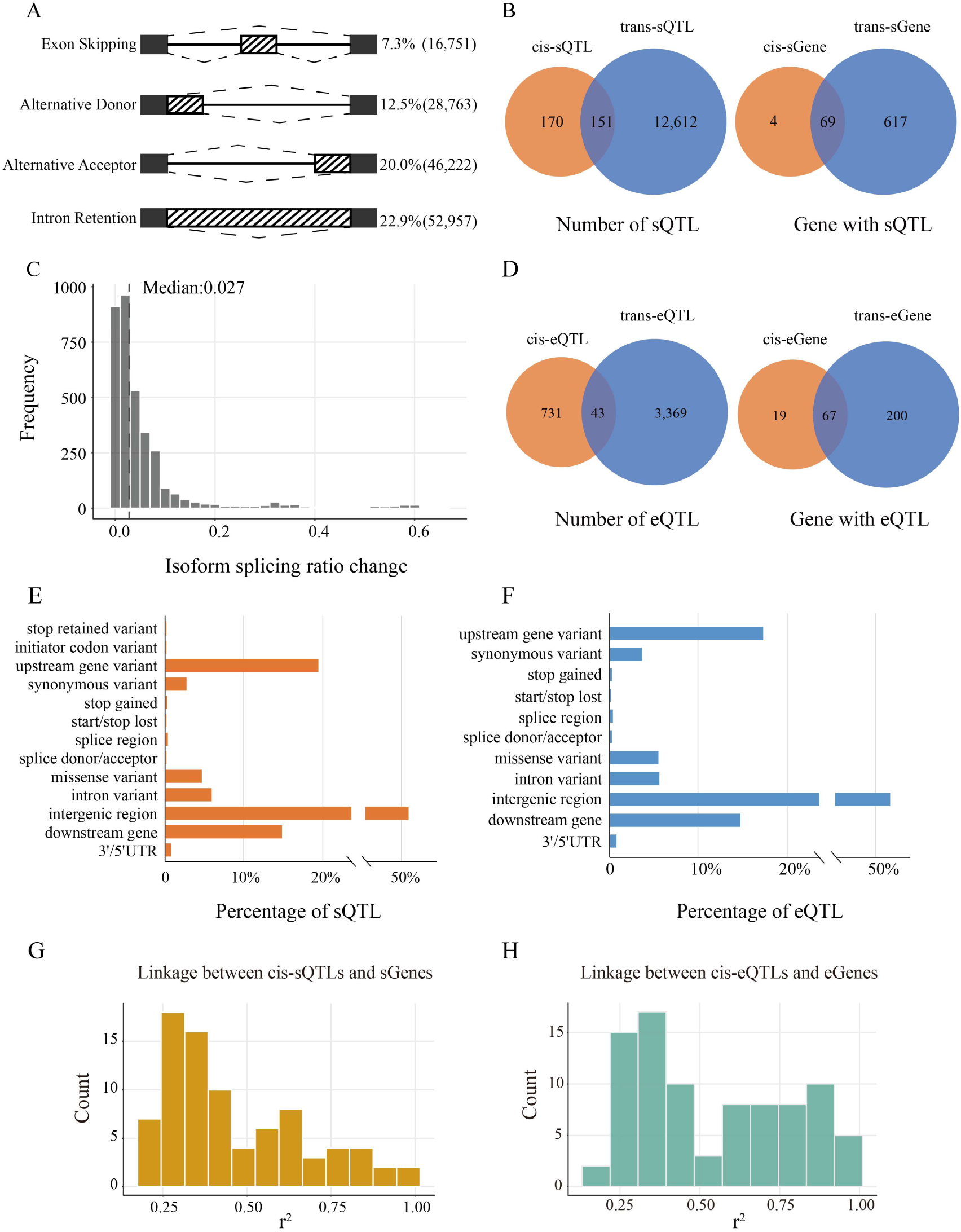
Statistic of the identified sQTL and eQTL. (A) Categorization of the AS events in assembly transcripts; **(B)** The distribution of cis/trans-sQTL and cis/trans-sGene; **(C)** The distribution frequency of alternative splicing ratio of these identified cis-sQTL; **(D)** The distribution of cis/trans-eQTL and cis/trans-eGene; **(E)** Genomic region annotation of cis-sQTLs; **(F)** Genomic region annotation of cis-eQTLs; **(G)** The distribution of linkage disequilibrium (r^2^) between cis-sQTL and its target sGene; **(H)** The distribution of linkage disequilibrium (r^2^) between cis-eQTL and its target eGene.

Most of the sQTLs and eQTLs were situated within the intergenic region, followed by the up/downstream gene regions (Figure 2E and 2F). The LD (r^2^) distribution of cis-sQTLs and cis-eQTLs showed an approximate left skewed pattern (Figure 2G and 2H). Furthermore, positional enrichment was conducted, and the majority of the cis-sQTLs (∼43.2%) and cis-eQTLs (∼80.4%) were distributed within 10 Mb intervals around the translation start site (TSS) with a slight downstream bias pattern related to their sGenes (sQTL-associated genes) and eGenes (eQTL-associated genes), respectively (Figure S5C and 5D).

The asymmetry of sQTLs and eQTL was also detected. A subgenome had 5,846 sQTLs (40 cis-sQTLs; 5,702 trans-sQTLs; 104 cis/trans-sQTLs), the B subgenome had 5,620 sQTLs (121 cis-sQTLs; 5,472 trans-sQTLs; 27 cis/trans-sQTLs) and the D subgenome contained only 1,467 sQTLs (9 cis-sQTLs; 1,438 trans-sQTLs; 20 cis/trans-sQTLs). Moreover, the A, B and D subgenomes contained 1,174 (121 cis-eQTLs; 1,031 trans-eQTLs), 1,501 (390 cis-eQTLs and 1,097 trans-eQTLs) and 1,468 (220 cis-eQTLs and 1,241 trans-eQTLs) eQTLs, with 22, 14 and 7 eQTLs shared by cis-and trans-eQTLs in the A, B and D subgenome, respectively. Taken together, the A subgenome tended to possess the most abundant sQTLs and eQTLs, followed by the B subgenome and D subgenome, suggesting an asymmetric evolutionary history among the three subgenomes in wheat.

### sQTLs involved in protein domain and miRNA targeting regulation

To understand the potential roles of these sQTLs, we predicted the conserved protein domains of sQTL-affected isoforms. In total, 259 (37.5%) sGenes were found to have domain changes in different isoforms, of which 244 genes showed domain gains/losses, and the other 15 genes showed domain changes (Table S6), suggesting that sQTLs may result in isoforms with different biological functions. For example, the trans-sQTL *AX-111055365* on chromosome 7A was found to control alternative splicing of *TraesCS1A02G253400* (P = 3.36E-08), resulting in its PEP conserved domain transformation into the PPDK domain in isoforms T3 (*TraesCS1A02G253400.3*) and T1 (*TraesCS1A02G253400.2*) (Figure 3A). Moreover, the transcript *TraesCS2D02G109900.2* was found to possess an acyl-CoA dehydrogenase domain, whereas the other splicing isoform *TraesCS2D02G109900.1* lost this domain, which is regulated by sQTL *AX-108885453* with the genotype GG showing a higher splicing rate than genotype AA, although the expression level remained consistent between them (Figure 3B). microRNAs (miRNAs) are a class of non-coding RNA that regulate gene expression by mediating targeted mRNA cleavage or translational inhibition [24]. Different isoforms cause mRNA sequence variations, which leads to changes in miRNA targeting sites. We analyzed the miRNA-binding sites of the sGenes and found that 277 (40.1%) genes had miRNA target site changes due to splicing. Among these genes, 47 sGenes exhibited gained/lost miRNA target sites, and 230 sGenes changed the target sites for different isoforms (Table S7). For example, cis-sQTL *AX-109620570* (chr6A_74974194, P = 9.01E-14) was associated with the sGene *TraesCS6A02G188100*, regulating it to produce three isoforms named T1 (*TraesCS6A02G188100.1*), T2 (*TraesCS6A02G188100.2*) and T3 (*TraesCS6A02G188100.3*) (Figure 3C). miRNA target prediction showed that the T2 isoform could be targeted by *tae-miR5049/1122c/1130b/1120b-3p*, whereas no miRNA target sites were found for isoforms T1 and T3. Furthermore, genotype analysis revealed that the AA haplotype showed a lower splicing rate and higher expression level, whereas the GG genotype showed high splicing with low expression. We postulated that sQTLs might not only affect the gene expression level, but also determine the gene isoform transcription and further shift the miRNA target binding sites, suggesting that sQTLs have vital effects on gene expression and function in wheat.

**Figure 3.**
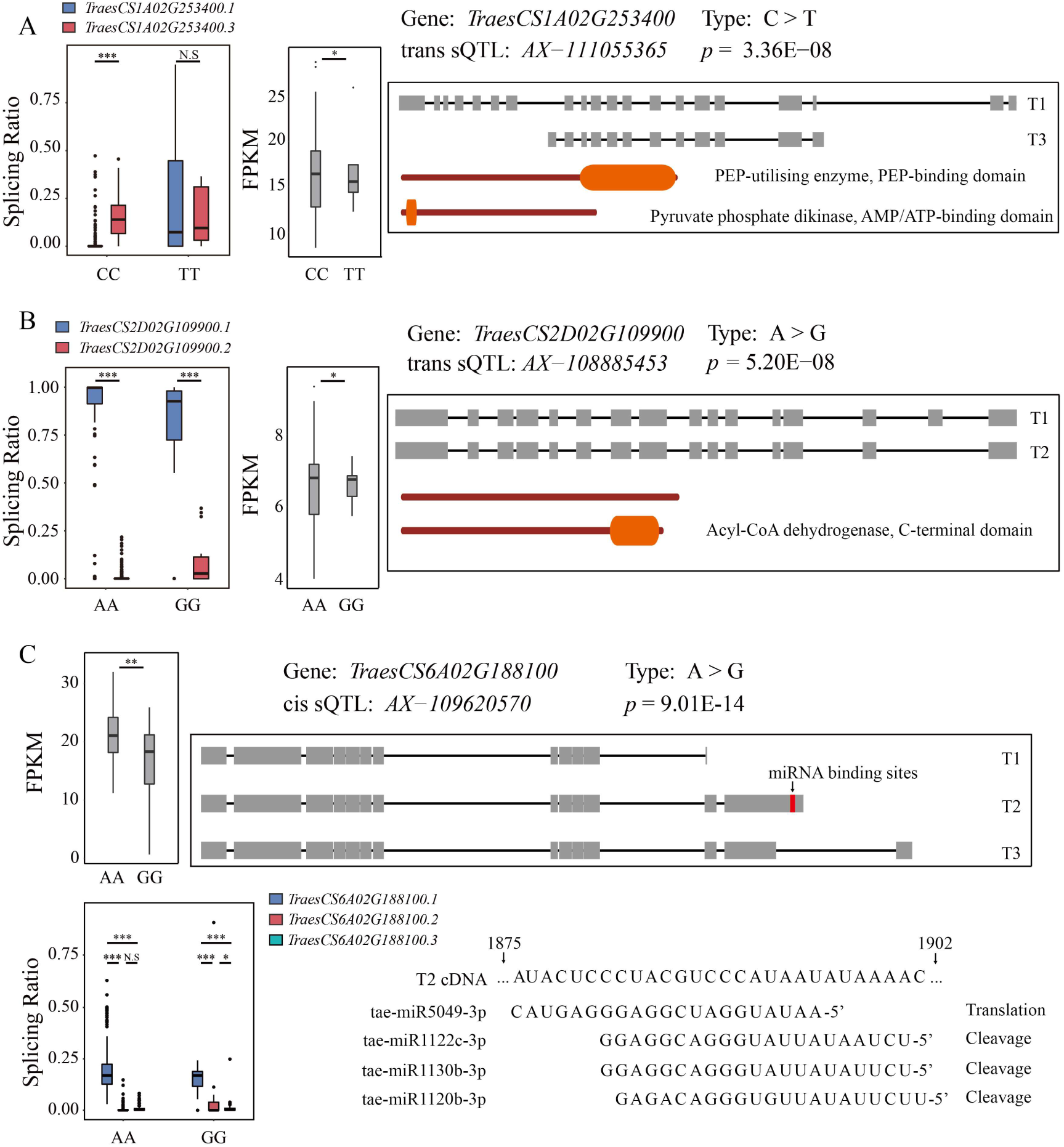
Genetic effects of sQTL regulating the target genes to display different protein domains and miRNA binding sites. (A) Significant trans-sQTL detected at *TraesCS1A02G253400*, showing differential isoforms has differential protein domains based on differential genotypes; **(B)** An example shows protein domain loss event in differential gene (*TraesCS2D02G109900*) transcript isoforms based on differential genotypes; **(C)** *TraesCS6A02G188100* was detected with a significant cis-sQTL, showing miRNA binding sites gained event in differential isoforms based on differential genotypes. *, *p*< 0.05; **, *p*< 0.01; ***, *p*< 0.001; N.S, not significant.

### Integration of phenotypic, genotypic, and transcriptomic signatures

Using a 11.396 Mb region as the boundary, we identified 3,154 co-localization events with sQTLs or eQTLs (Figure 1; Table S8). Among these loci, 67 SNPs were defined as cis-sQTLs and cis-eQTLs simultaneously, accounting for 20.87% and 8.66% of the total members, respectively (Figure 4A). Correlations between GWAS signals and cis-sQTLs/eQTLs were further determined. There were 6,488 co-localization events between 738 eQTL and 408 GWAS signals. Otherwise, 1,947 co-localization events were identified between 257 sQTL and 599 GWAS signals (Table S9-11). Consistent with a previous study [20], only a tiny portion of the genes (3, 1.92%) were found to be shared by sGenes and eGenes. Therefore, the sQTLs and eQTLs that appeared in the co-expression network attract extensive attention as they regulate mRNA level variation and AS variation in gene sets with the co-expression patterns.

**Figure 4.**
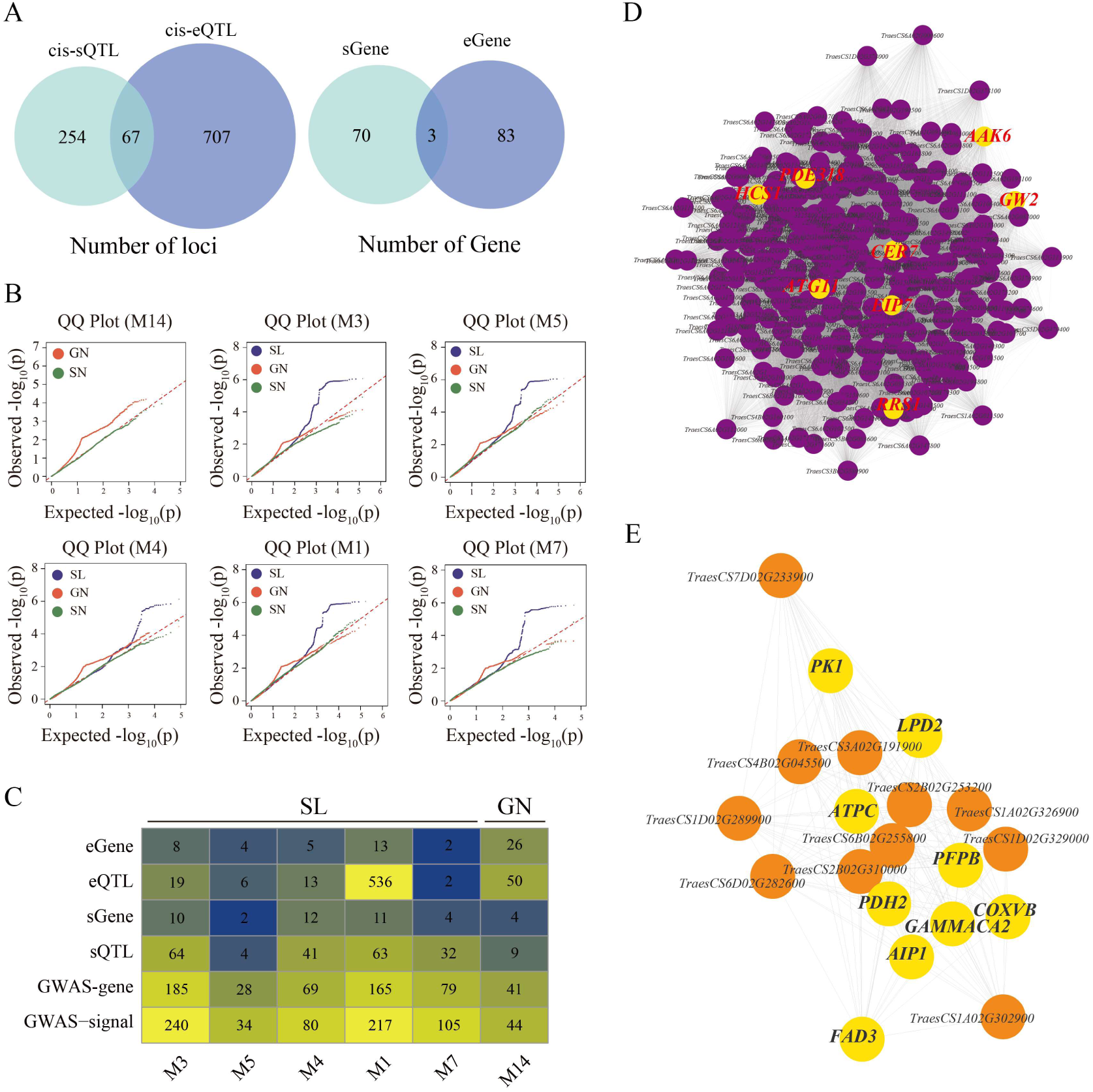
Integration of the eQTL and sQTL with the co-expression networks of wheat spike development. **(A)** Overlap between cis-sQTL and eQTL, sGene and eGene; **(B)** QQ plot of GWAS signals of target traits corresponding to different co-expression modules. For each module, the trait with the most significant signals from the expected value is the optimal trait regulated by the module; **(C)** The number distribution of each variant type and target gene in each related module; **(D)** Co-expression network (M14) significantly (P = 2.13E-12) associated with GN; **(E)** Co-expression network (M1) significantly (P = 5.92E-03) associated with SL.

Based on the co-expression modules identified, the potential relationships between sGene/eGene and the spike-related traits were further investigated. GWAS analysis identified 859 over-represented signals with SL, 611 with GN and 263 with SN, as well as 1,498 with grain-related traits (Table S1). We enriched signals of spike-related traits with a QQ plot, and the number of signal points significantly related to traits was taken as the standard (Figure 4B). Consequently, 6 trait-related WGCNA modules were detected and verified by a hypergeometric test, of which M14 module controlled GN, and SL was regulated by five modules (M1, M3, M4, M5 and M7) (Figure 4C; Figure S6-9). We further integrated PO, TO and KEGG results and found that these six modules were all related to plant development. In particular, PO terms, including plant ovary, glume and ovule development stages, were significantly enriched in the M1 module. Moreover, KEGG analysis found that they were enriched in the autophagy pathway (Table S12). Subsequently, a series of spike development genes was found in these modules. For example, *TraesCS6A02G189300*, the hub gene in the GN-related module, is the ortholog of *OsGW2* that controls grain size in rice [25]. In addition, *TraesCS6B02G279300*, the hub gene in the M1 module related to SL, is the orthologous gene of *AtYDA* that regulates coordinated local cell proliferation and further shape the morphology of plant organs through the MAPK cascade pathway [26].

### Molecular module of grain number of per spike (GN)

A total of 284 genes were present in the GN-related module, including 26 eGenes and 4 sGenes. Obviously, eQTLs and sQTLs have relatively independent regulation patterns for this trait. The expression of five genes (*TraesCS6A02G088700*, *TraesCS6A02G153400*, *TraesCS6A02G170900*, *TraesCS6A02G181600*, and *TraesCS6A02G186300*), was regulated by the same eQTL locus, *AX-108754757,* on chromosome 6A (Figure 5A). Furthermore, *AX-108754757* also showed a strong association with GN through GWAS analysis (Figure 5B). Based on this eQTL, all of the wheat varieties were divided into two groups, with the TT (85) or CC (4) genotype, respectively. Notably, varieties with the TT genotype showed lower expression levels but more grain number per spike than those with the CC genotype, suggesting that GN was controlled by this eQTL through negatively regulating the expression of the target genes (Figure 5C and 5D). From the co-expression network, we found that these five genes formed the hub of this module, playing a key role in regulating other genes in this module to achieve its function (Figure 5E). Moreover, the other 279 genes in this module also showed negative associations between the expression level and grain number (Figure 5F and 5G). These results suggested that the M14 module strongly correlated with the GN trait with a negative effect, especially the eQTL locus *AX-108754757*, which is a vital regulator that controls the expression of five hub genes.

**Figure 5.**
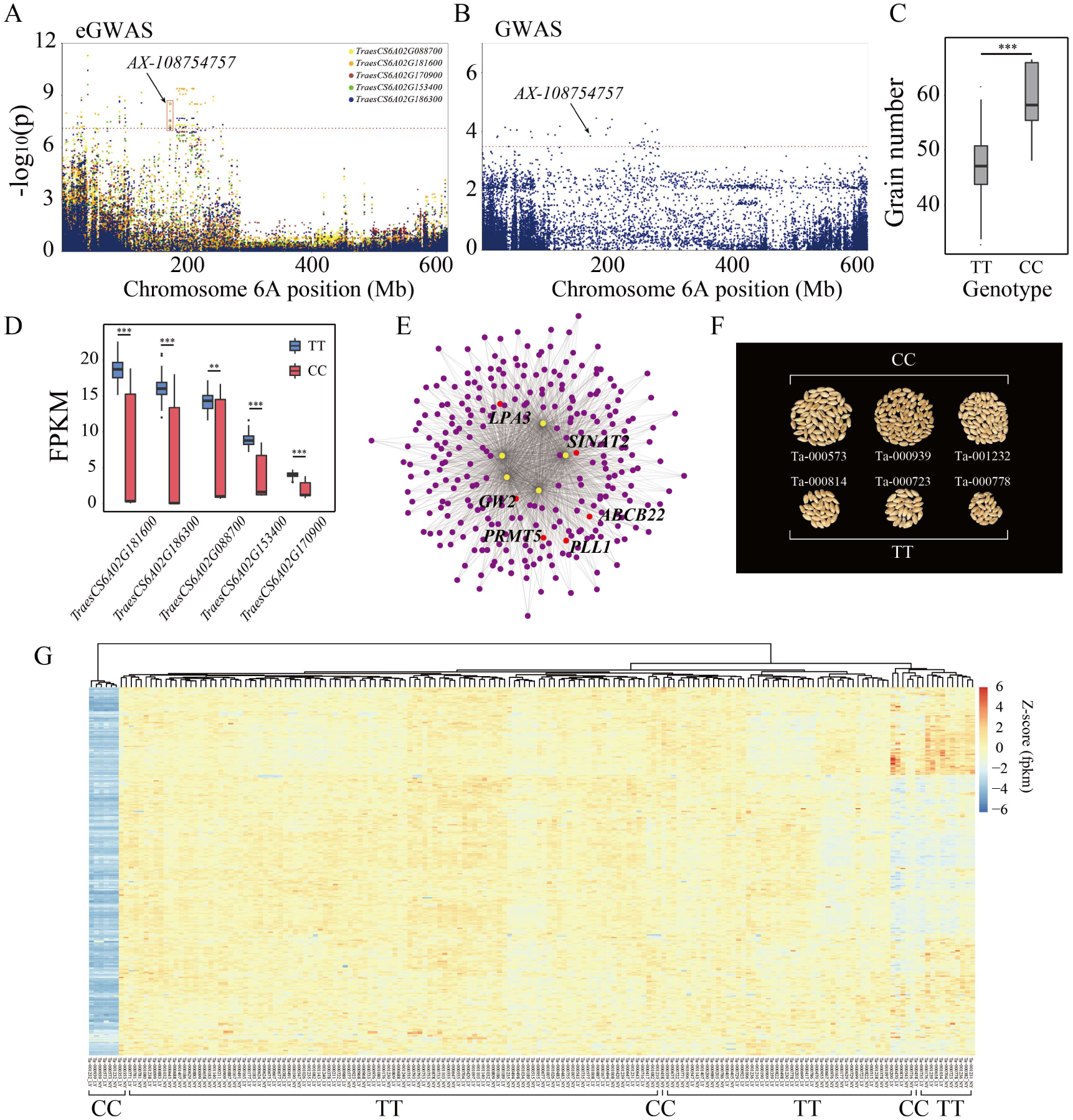
Co-expression module associated with grain number per spike (GN). (A) Manhattan plot of eGWAS between SNP and five genes expression level. Red horizontal dashed line indicates the genome-wide significance threshold (–log_10_P = 7.1); (B) Manhattan plot of GWAS between SNP and GN. Red horizontal dashed line indicates the genome-wide significance threshold (–log_10_P = 3.5); (C) Effects of different genotypes on phenotypic traits; (D) Effects of different genotypes on five genes expression level; (E) Five significant eGenes in M14 module were highly connected by other co-expression genes and located in hub position; (F) Phenotype of GN in different genotype; (G) The expression heatmap shows normalization FPKM of co-expression genes were affected by common eQTL of five hub genes. *, *p*< 0.05; **, *p*< 0.01; ***, *p*< 0.001; N.S, not significant.

Compared with eQTLs, sQTLs affect gene expression by regulating AS events. Two linked SNPs (r^2^ = 0.761) on chromosome 6A (*AX-94670617* and *AX-110511094*) were associated with the alternative splicing of *TraesCS6A02G047900* in this module (Figure 6A), of which locus *AX-94670617* (P = 6.91E-04) also associated with grain number by GWAS and *AX-110511094* (P(sQTL) = 2.78E-09; P(eQTL) = 2.60E-08) and was found to regulate AS events and gene expression levels (Figure 6B). The haplotypes were further analyzed using the two linked loci, and four main haplotypes were obtained (Figure 6C). Compared with other haplotypes, hap2 showed the lowest alternative splicing ratio but had the highest grain number per spike (Figure 6D–F). Meanwhile, *TraesCS6A02G047900* was also highly linked to the five hub eGenes (Figure 6G), suggesting that sQTL also had a regulatory effect on GN.

**Figure 6.**
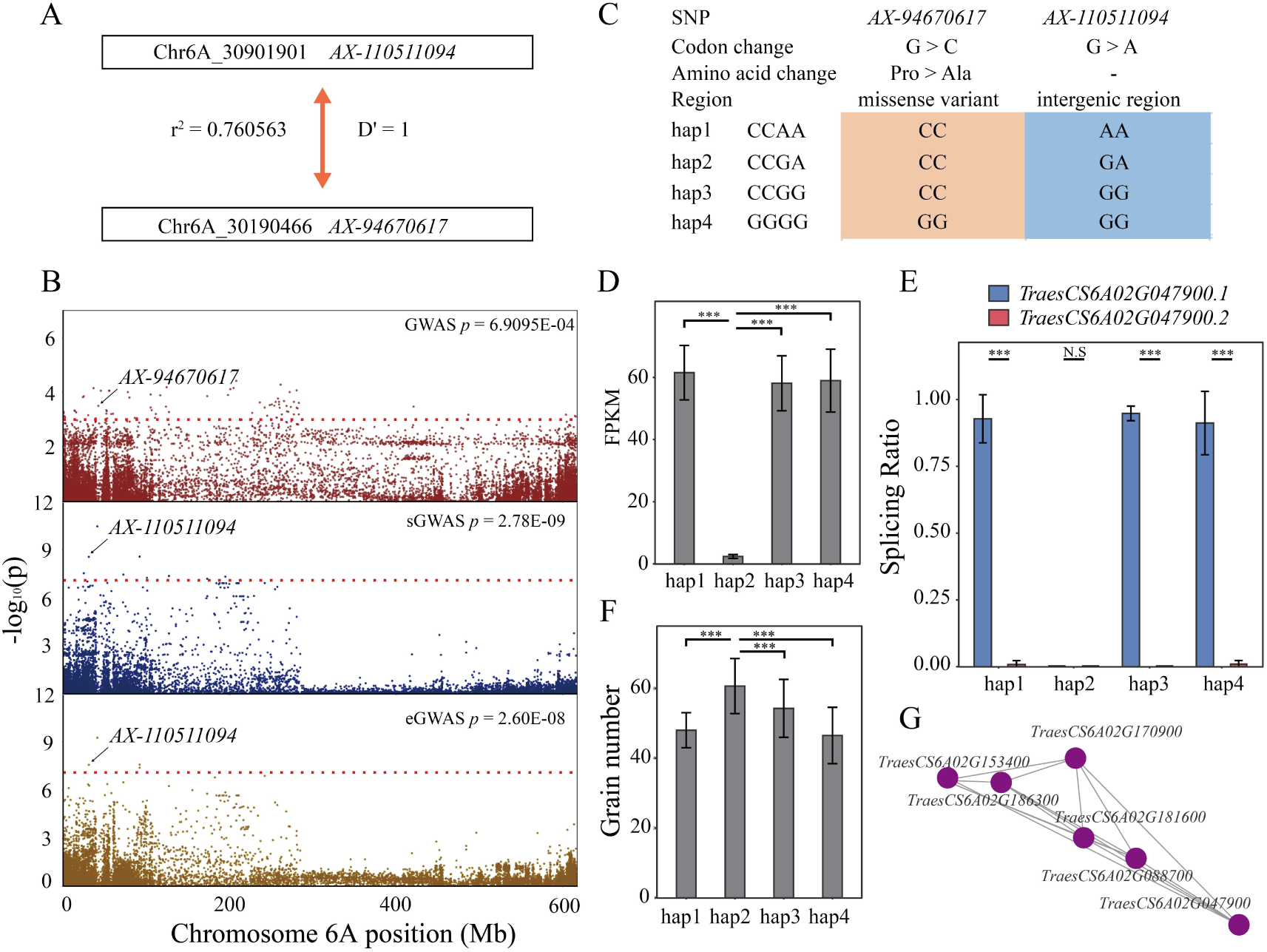
*TraesCS6A02G047900* is the hub gene in GN-related module and its expression and splicing were regulated by locus *AX-110511094*. (A) Linkage disequilibrium (r^2^, D’) was calculated between s/eQTL and GWAS signal of *TraesCS6A02G047900;* (B) Manhattan plots of GWAS results in phenotype variation, gene expression variation and alternative splicing variation. sQTL and eQTL were the same SNP and GWAS signal was another SNP. Red horizontal dashed line indicates the genome-wide significance threshold (s/eGWAS –log_10_P = 7.1; GWAS –log10P = 3.5); **(C)** Haplotype genotype between s/eQTL and GWAS signal was constructed and the main haplotypes were screened and analyzed; **(D)** Effects of different haplotypes on gene expression level; **(E)** Effects of different haplotypes on alternative splicing ratio; **(F)** Effects of different haplotypes on phenotype value; **(G)** Co-expression network of *TraesCS6A02G047900* and other hub genes. *, *p*< 0.05; **, *p*< 0.01; ***, *p*< 0.001; N.S, not significant.

### Molecular modules of spike length (SL)

We found 5 modules associated with spike length, and most of the eGenes and sGenes were detected in the M1 module. *AX-111592099* was found to be associated with the AS ratio and expression level of *TraesCS7B02G442100* (Figure 7A). The AS ratio of its transcript T1 (*TraesCS7B02G442100.1*) was significantly higher than the alternative splicing ratio of T2 (*TraesCS7B02G442100.2*) under the regulation of genotype GG (N = 57). In contrast, the AS ratio of T2 increased significantly under the genotype AA (N = 32), although the AS ratio of T1 was still higher than the AS ratio of T2 in general (Figure 7B). The expression level of *TraesCS7B02G442100* controlled by eQTL *AX-111592099* was also significantly different, with a high expression level in the GG genotype and low expression in the AA genotype, showing that the expression level was mainly based on T1, and the increase in T2 may decrease the expression level (Figure 7C). Based on the association results, we found that sGWAS and eGWAS have the same two highly linked blocks with a size of 120 kb and 420 kb, respectively (Figure 7D). 14 genes were found in these regions, and their orthologues showed significant biological functions (Figure 7E). Among them, *ENSRNA050019690*, orthologues to *U1 SnRNA*, was found to be involved in the splicing of pre-mRNA, confirming that sQTL can regulate splicing variations. *TraesCS7B02G442100* is the orthologue of *SUPPRESSOR OF DRY2 DEFECTS1 (SUD1)*, which encodes a putative E3 ubiquitin ligase and regulates 3-hydroxy-3-methylglutaryl coenzyme A reductase activity without apparent changes in protein content [27]. In the co-expression network, 9 genes were highly linked to *TraesCS7B02G442100* (Figure 7F). The expression levels of these genes also displayed significant differences between the two genotypes, of which 5 genes tended to be significantly up-regulated from GG to AA, while the other genes tended to be significantly down regulated from AA to GG (Figure 7G). Interestingly, the orthologues of these highly linked genes were always related to protein modification, especially ubiquitination. These include *TraesCS7B02G437600* (*PBC1*), *TraesCS7D02G510600* (*PBC1*), *TraesCS5A02G557300* (*ALPHAC-AD*), *TraesCS4B02G395700* (*ALPHAC-AD*) [28], *TraesCS7B02G436500* (*CHMP1A*), *TraesCS7A02G524600* (*CHMP1A*) [29, 30] and *TraesCS7B02G442100* (*SDU1)*.

**Figure 7.**
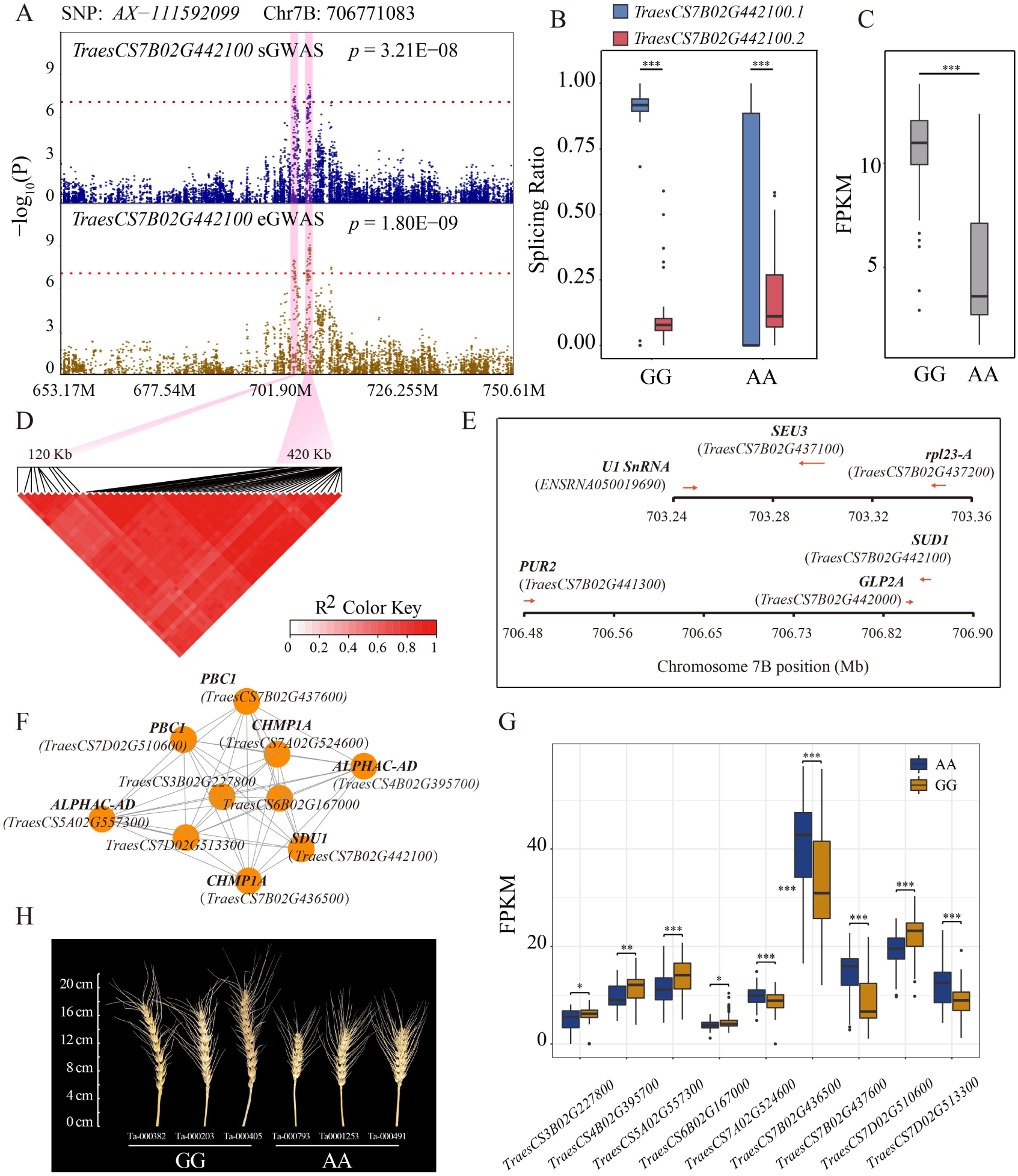
Co-expression module associated with spike length (SL). (A) Manhattan plot of eQTL and sQTL loci in *TraesCS7B02G442100*. Red horizontal dashed line indicates the significance threshold (–log_10_P = 7.1); **(B)** Effects of different genotypes on gene expression level; **(C)** Effects of different genotypes on alternative splicing ratio; **(D)** Two high-linkage LD blocks were constructed in sGWAS and eGWAS, and both GWAS are highly coincident; **(E)** Orthologous genes in the LD blocks; **(F)** Co-expression network of *TraesCS7B02G442100;* **(G)** The expression levels of 9 genes in the network were also regulated by the same SNP; **(H)** Phenotype of SL in different genotype. *, *p*< 0.05; **, *p*< 0.01; ***, *p*< 0.001; N.S, not significant.

Notably, these genes were related to protein complexes, such as E3 ubiquitin ligase, ESCRT-dependent multivesicular body (MVB) formation and autophagosomes [30]. These results showed that the gene expression level and AS ratio of *TraesCS7B02G442100* were controlled by genetic factors, and highly linked genes with the same expression pattern and function in the module.

To evaluate the relationship between the module and the actual phenotype, 2 out of 9 highly linked genes were found to have GWAS signals with SL in gene exon regions (Figure S10A and 10B). In *TraesCS4B02G395700*, *AX-108983060* (chr4B_670404267, P = 4.83E-04) and *AX-110916949* (chr4B_670409830, P = 8.05E-05) located in the 9th exon and 27th exon, respectively (Figure S10C and 10D). *AX-108983060* was a synonymous variant, and *AX-110916949* changed the amino acid sequence from Gln to Glu (Figure S10E). In *TraesCS4B02G557300*, *AX-110442752* (chr5A_708411623, P = 1.31E-04) and *AX-111171087* (chr5A_708417656, P = 6.64E-04) located in the 2nd exon and 20th exon, respectively, and both SNPs caused synonymous variants (Figure S10F). The wheat population had three main haplotypes in this gene, of which hap3 had the largest spike length, followed by hap2 (Figure S10G). Ranking to haplotypes, hap4 had the largest spike length, followed by hap3 and hap1 (Figure S10H). These results confirmed that the sQTLs and eQTLs in M1 module had regulatory effects on wheat spike length, and a set of genes that related to spike length was identified.

Additionally, the SNP locus *AX-109363523* regulated the variation in expression of T*raesCS7A02G279100* as well as the splicing variation of *TraesCS7A02G283100* (Figure 8A–D), meaning that it functioned as both an eQTL and a sQTL. Furthermore, *TraesCS7A02G279100* present in the M1 module, and *TraesCS7A02G283100* was found in the M7 module. In co-expression module, 28 genes were highly linked with *TraesCS7A02G283100*, and 20 genes were highly linked with *TraesCS7A02G279100.* Furthermore*, AX-109363523* also regulated the variation in expression of the highly linked genes in each module (Figure 8E–H). This result suggested that the same genetic variation could have different effects on spike development by mediating different molecular modules through regulating gene expression and splicing. We also found that in the same module (M1 module), the expression level of *TraesCS5D02G454700* was regulated by *AX-111930824* (eQTL, P = 5.90E-08), and the splicing variation of *TraesCS5A02G547700* was regulated by *AX-109874238* (sQTL, P = 3.0E-08) (Figure S11A-D). In addition, 20 genes were highly linked with *TraesCS5D02G454700* and *TraesCS5A02G547700* (Figure S11E), of which 7 displayed differential expression between the two genotypes (Figure S11F). These results further demonstrated the complexity of the genetic network underlying spike length in wheat.

**Figure 8.**
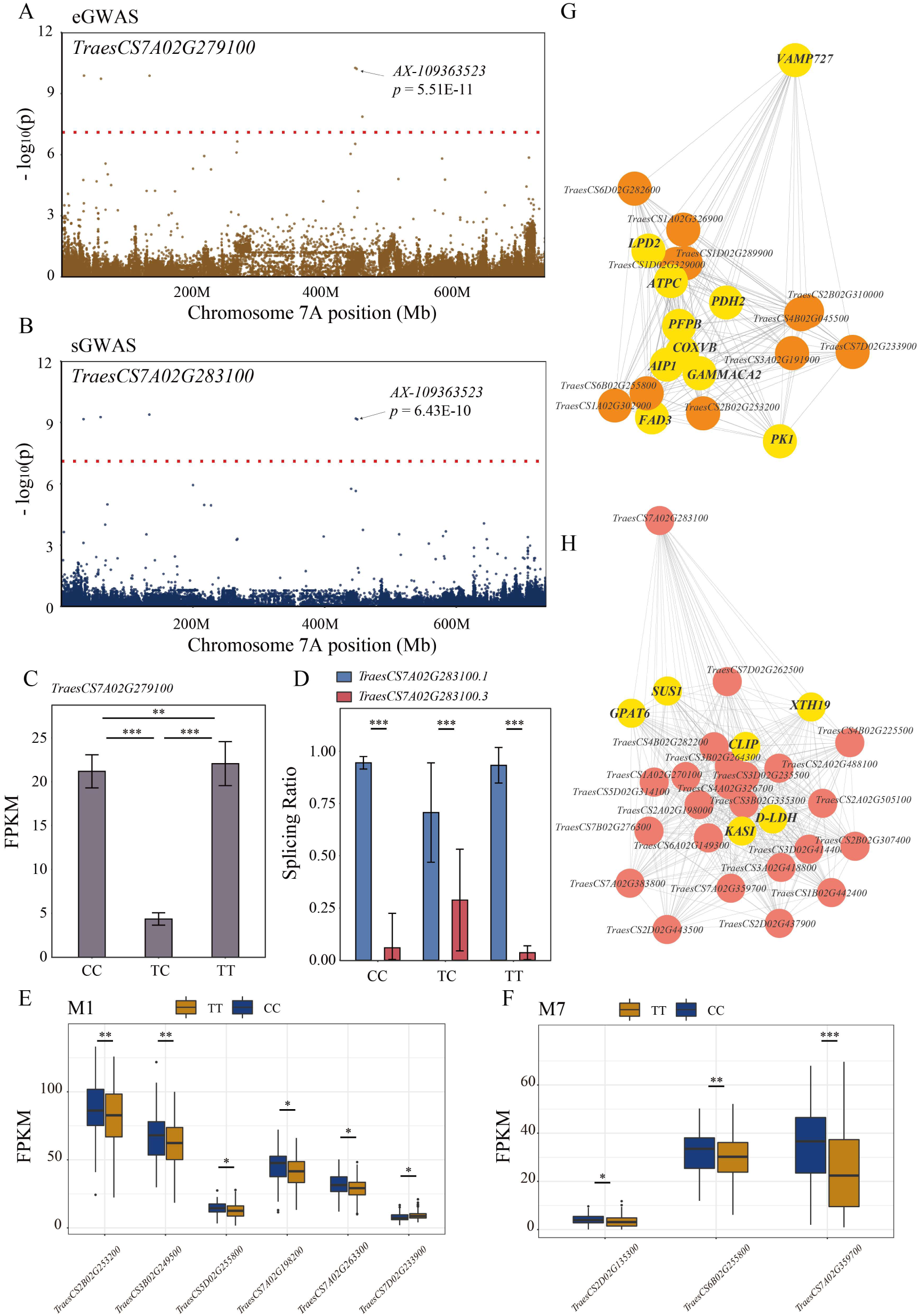
One SNP locus *AX-109363523* regulated the expression variation of *TraesCS7A02G279100* and the splicing variation of *TraesCS7A02G283100*. (A) Manhattan plot of eGWAS shows the relationship between the SNP locus *AX-109363523* and gene expression level (FPKM value) of *TraesCS7A02G279100*; (B) Manhattan plot of sGWAS shows the relationship between the SNP locus *AX-109363523* and the splicing (AS Ratio) of *TraesCS7A02G283100*; (C) Gene expression variations among different genotypes; (D) Alternative splicing variation among different genotypes; (E) Expression variation of co-expression genes highly linked with *TraesCS7A02G279100* in M1 module; (F) Expression variation of co-expression genes highly linked with *TraesCS7A02G283100* in M7 module; (G) Co-expression network of *TraesCS7A02G279100*; (H) Co-expression network of *TraesCS7A02G283100*. *, *p*< 0.05; **, *p*< 0.01; ***, *p*< 0.001; N.S, not significant.

## Discussion

As the world’s most widely grown crop, wheat plays an important role in meeting the challenges of food security under changing global climate [1]. In the past decade, a slower and slower growth rate of yield per unit area has occurred in China and other main wheat-producing countries, indicating a bottleneck in yield potential due to conventional hybridization breeding methods [31]. It is well known that wheat yield is mainly determined by spike-related traits. Thus, dissection of the causal genes and the mechanism underlying spike-related traits is crucial for wheat yield improvement.

Although some genes controlling spike architecture have been reported in rice, maize and barley [32], it remains challenging to identify genes conferring spike-related traits in wheat due to these being complicated quantitative traits and the complex wheat genome [33]. In this study, we systematically investigated the genetic basis of wheat spike architecture from three perspectives: genetic variations, expression variations and splicing variations. From the perspective of genetic variations, 3,231 loci were identified to be associated with 12 spike-related traits based on GWAS analysis. The genome-wide LD decay (r^2^ = 0.2) of the wheat population was calculated to be 11.396 Mb based on 590,136 SNPs, and the genes in the LD interval with GWAS signals were considered to be potential trait-related genes. Although we obtained some putative genes related to spike architecture, the key causal genes and their regulation modules were not well determined. The phenotypic diversity of organisms results directly from the regulation of gene expression rather than genetic variations [34]. To better understand the regulatory mechanism of spike architecture, we quantified the mRNA abundance and splicing through population RNA-seq analysis and then eQTLs and sQTLs were identified. In total, 4,143 eQTLs and 12,933 sQTLs were obtained, representing the first expression variation and splicing variation landscape in wheat, especially for spike-related traits. Then, a co-expression network was constructed based on the 38,285 expressed genes in 89 varieties to obtain the module and gene sets associated with wheat spike architecture. Functional enrichment analysis of co-expression modules showed that 10 modules were significantly enriched in some terms related to spike and grain development, including ubiquitination process, plant ovary development and glume development. In maize, Liu *et al* used Mendelian randomization methods that adopted a two-step least squares (2SLS) to link eQTL and traits through estimating the effect of x (normalized gene expression) on y (traits) to reveal the relationship between gene expression levels and phenotypic traits, which provided a more reasonable estimation of the influence of gene expression on trait performance [18]. Our study tried to integrate the GWAS signal, eQTLs and sQTLs based on a co-expression network, which could not only identify the key genes controlling the traits but also obtain their molecular modules, which is vital for molecular breeding. Although the MR method was not used, the key genes related to spike architecture were also found by the association of genetic variations with agronomic traits, gene expression and the AS ratio, and the reliability was verified by phenotype evaluation.

We found that one module related to GN and 5 modules related to SL in this study. Furthermore, 50 cis-eQTLs associated with the expression of 26 genes and 9 cis-sQTLs associated with the AS ratio of 4 genes were further identified in the GN module, while 576 cis-eQTLs and 204 cis-sQTLs were identified in five SL modules, suggesting that SL seems more complex and to be regulated by many genes and patterns. The number of gene expression variations and AS variations in the gene co-expression module showed that the number of eQTLs was greater than the number of sQTLs, and eGene was greater than sGene. Gene expression variation contributed more to the regulation of spike-related traits, while the variation in AS had stronger effects on the conserved domains of some hub genes and miRNA binding sites. Subsequently, we found that only a small number of SNPs could affect both gene expression and AS, which was consistent with a previous study in maize [20]. Moreover, GWAS signals were integrated, and we identified 378 unique and 44 unique signals in the SL and GN modules respectively, with 290 and 41 unique signal-related genes (Table S13, 14). GWAS signals in these modules further verified the correlation between the modules and the target traits and they could also be regulated by eQTLs and sQTLs to form sophisticated and precise regulatory modules to control complex traits.

In the GN-related module, we found that the expression levels of five hub genes were regulated by *AX-108754757* to negatively correlate with GN. In this module, the *OsGW2* orthologue, *TraesCS6A02G189300,* was found, and this gene acts as an E3 ubiquitin ligase to regulate grain width, grain weight, panicles per plant and grains per panicle [35]. Moreover, a gene set that was highly correlated with *GW2* was found to form a regulation module, which lays the foundation for genetic improvement of grain number. In the SL modules, we found a series of genes related to protein modification and ubiquitination processes, of which *TraesCS7B02G442100* was regulated by both eQTLs and sQTLs. Co-expression gene sets analysis showed that 9 genes were connected with *TraesCS7B02G442100* and the expression levels of 9 genes were also regulated by *AX-111592099*. With genotype GG to AA, the expression levels of 4 genes showed positive changes and 5 genes showed negative changes. Among these genes, six genes had orthologues with Arabidopsis, including *PBC1*, *ALPHAC-AD*, *CHMP1A* and *SUD1*. Notably, the function of these six genes was related to protein transport and synthesis, especially the E3 ubiquitin ligase pathway and autophagy pathway. At the phenotypic level, three samples were randomly selected for measurement. The average spike length of the GG genotype was approximately 18 cm, the average spike length of the AA was 14 cm, which verified the contribution of the variation in *AX-111592099* to spike length (Figure 7H).

## Conclusion

In this study, we reported a comprehensive set of eQTLs and sQTLs mapped in the co-expression modules of spike architecture in wheat. Our analysis revealed that gene expression variations and splicing variations contributed significantly to wheat spike-related traits. By integrating eQTL, sQTL and GWAS signals as well as co-expression networks, one module associated with GN and five modules associated with spike length were obtained and the hub causal genes were also identified. We also demonstrated that the ubiquitin ligase pathway is strongly involved in wheat spike development and is affected by multiple patterns of gene expression variation and AS variation. This study lays the foundation to better understand the genetic basis underlying spike architecture in wheat and sheds light on dissecting the molecular mechanism of complex agronomic traits through associative transcriptomic methods.

## Methods

### Plant materials and growth conditions

A total of 89 representative wheat varieties selected from 5,000 accessions worldwide [23], which were grown in the Nanyang (NY) and Luoyang (LY) trial, China in the 2018-2019 cropping season, were used in this study. At the heading stage, three spikes from different individuals of each accession were collected and then bulked together for subsequent experiments and RNA sequencing. All samples were collected from 9 to 11 a.m. in the field.

### RNA-sequencing data mapping and transcript quantification

Total RNA was isolated using the Qiagen Plant Tissue RNA Isolation kit following the instructor’s protocol. RNA-seq was performed using Illumina® HiSeq X Ten platform with paired-end 150 bp. The raw RNA-seq reads were subjected to FastQC v0.11.7 and Trimmomatic v0.36 toolkit to remove the adapter, and filter the low-quality reads as well as the contamination of unknown nucleotides [36]. The obtained clean reads were aligned onto the IWGSC RefSeq (version 1.1) for each sample using HISAT2 v2.1.0 [37]. For the mapping bam files, the SAMTools v1.3 was used to filter the reads with mapping quality lower than 60 [38]. Furthermore, the StringTie v2.0.4 was employed to assemble the transcripts for each accession initially and then combined them together into the unified dataset of transcripts using the ‘merge’ toolkit [39,40]. Known and novel transcripts were quantified for each accession using StringTie software with default parameters, and the transcripts shared by LY and NY were retained for further analysis. Notably, novel transcripts located within the intergenic region were excluded from our analysis. Ultimately, a comprehensive pipeline was used to determine the expression threshold of transcriptional isoforms’ FPKM (transcriptional fragments per kilobase in the per million mapping reads) (Figure S12) as previous study described [41]. The transcripts with FPKM lower than 0.52 with representation lower than 5% varieties were abandoned.

### Co-expression network analysis

The following filtration was employed for WGCNA analysis [42]. For genes, the FPKM > 0.52 with representation in > 5% of the total varieties and the variance of expression > 5 was retained [20]; for samples, the standardized sample network connection Z score should be lower than 2.5. The optimal soft threshold power was simulated with an integer number ranging from 1 to 10 using pickSoftThreshold toolkit embedded in WGCNA. The suitable power was set at 7 by calculating the scale-free topology (r^2^ > 0.85) fitting index with various power values. Based on the predefined soft threshold power, the adjacency matrix was constructed and was then converted into a topological overlap matrix as an input for hierarchical cluster analysis. Subsequently, the topological overlap matrices were clustered hierarchically using average linkage hierarchical clustering with the ‘1-TOM’ option as a dissimilarity measure. Then, the cutreeHybrid toolkit in the R package dynamicTreeCut was used to determine the network modules that existed in the Normal dataset (used here as a reference data set) with the following parameters: a relatively large minimum module size (minClusterSize) = 50, merge threshold of 0.25, and a medium sensitivity(deepSplit) = 2, other parameters set as default. Remarkably, genes within the grey module means no assignment, that is no co-expressed genes were found for the genes within this module. Using the signedKME toolkit, the distance between gene expression level and eigen gene of specified module was measured to quantify the closeness between the gene and the module, and further to detect the hub genes within the module. The modules are numbered M1-M25 in descending order of gene number. The genes with the highest module member score were considered as the hub gene within the corresponding module.

### Genome-wide association study analysis

Genotype data of these 89 wheat varieties was performed using the wheat 660K genotyping assay by Beijing CapitalBio Technology Company (http://www.capitalbiotech.com). SNP genotype calling and allele clustering were processed with the polyploid version of Affymetrix Genotyping ConsoleTM (GTC) software. The physical positions of the SNPs were obtained from the Triticeae Multi-omics Center website (http://202.194.139.32/). Then, 12 spike-related traits, including spike number per plant (SN), grain number per spike (GN), spike length (SL) and other 9 grain-related traits (GA, GR, GC, GD, GL/GC, GW/GC, GL/GW, GA/GC, TGW) were used for performing GWAS analysis. Considering to the environment representations, the arithmetic mean values were calculated for each trait. Based on the compressed MLM model, GWAS was conducted using TASSEL v5.0 [43]. For the compressed MLM model, the following equation was employed in our study:

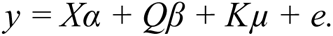

In these equations, *y* represents phenotype, *X* represents genotype, *Q* is the PCA matrix instead of the *Q* matrix and *K* is the relative kinship matrix. *Xα* and *Qβ* represent fixed effects, and *Kμ* and *e* represent random effects. The top three principal components were used to build up the *Q* matrix for population structure correction. The matrix of simple matching coefficients was used to build up the *K* matrix, and this step was followed by compression. Genotype (590,136 SNP quality-controlled loci) and environment (one years in two locations) were treated as random effects in the compressed MLM to estimate the best linear unbiased predictions (BLUPs).

### eQTL and sQTL analysis

For eQTL GWAS, the expression level of each gene was defined as the phenotype. For sQTL GWAS, only genes with more than two transcripts were used. By taking the total transcriptional abundance of each gene as the denominator, the splicing rate of each transcript isoform (subtype) was calculated and defined as the phenotype for association analysis [20]. The GWAS of eQTL and sQTL were also performed based on the compressed MLM model by incorporating the kinship coefficients, population structure, and hidden confounding factors using the TASSEL v5.0. The command option was *‘-mlm-mlm VarCompEst P3D-mlmCompressionLevel Optimum’*.

Furthermore, a Bonferroni-corrected (P-value <= 0.05/N (Number of SNP), N=590,136) test was conducted to calibrate the associated confidence with α < 0.05. The relative chromosome position of eQTL/sQTL and their associated genes’ TSSs were determined based on the genome annotation file. SNP and its target gene located on the same chromosome with r^2^ > 0.2 were considered as cis-eQTL/sQTL, otherwise, the eQTL/sQTL was treated as a trans-eQTL/sQTL.

### Overlap of cis-eQTL and cis-sQTL

If the signals of sQTL-GWAS, eQTL-GWAS and QTL-GWAS co-located together, this region was predicted to be with strong confidence that the SNPs and target genes are significantly relevant to the trait through the genetic regulation of gene expression or alternative splice. According to the linkage disequilibrium analysis, the linked region length was set at 11.396 MB for sQTL, eQTL and GWAS signals. The Circos tool v0.69-8 was used to display the overlap regions among these three types of association signals [44].

### Identification of spike trait related modules

In order to identify the most relevant modules of spike traits, we refer to the principle of module trade correlation in WGCNA. Based on LD (r^2^ > 0.2) and on the same chromosome, eGene and GWAS gene with linkage effect in each module were screened [45]. Then, we screened the corresponding GWAS signal loci as the basis to evaluate whether the module is related to spike traits. To assess significance, all signals of each module of each phenotype were filtered statistically by QQ-plot analysis [15]. The most relevant trait in each module was defined as the trait whose P-value exceeds the most observed value. The phyper function in R was used for hypergeometry test.

### Gene function enrichment analysis

Gene Ontology (GO) [46], Plant Ontology (PO), Plant Trait Ontology (TO) and KEGG analysis was carried out to evaluate the potential functions and metabolic pathways for each WGCNA module genes [47, 48]. The over-represented GO terms were determined using the online toolkit AgriGO v2.0 [49] with the following criteria: at least five gens; false discovery rate (FDR) <0.05. In order to determine the GO/TO terms of wheat genes, the BLAST search was conducted between the wheat genes and the GO/PO database, which was downloaded from the Plant Ontology Consortium (POC) (www.plantontology.org). The PO and TO enrichment were performed using the ClusterProfiler package of R v3.6.2. KOBAS v3.0 was used for KEGG pathways enrichment [50].

### miRNA target and conserved domain analysis

To determine the candidate miRNA targets affected by sQTL regulation, the online psRNATarget (http://plantgrn.noble.org/psRNATarget) [51] was used to predict the potential binding sites by sQTL using all wheat mature miRNA available from miRBase 22 (http://www.mirbase.org/). The Hidden Markov Model (HMM) of 18,197 conserved domains in the PFAM database (http://pfam.xfam.org/) was used as query to search against the known and novel transcripts using the HMMER v3.3.1 tool with E-value < 1E-5 as threshold, the domain hit results of novel and known transcripts were compared in a pairwise manner [52].

### Orthologous gene analysis

In order to better clarify the potential function of the identified key genes, the functionally validated genes in *Arabidopsis* and rice were downloaded from the TAIR (https://www.arabidopsis.org/) and Ricedata (https://www.ricedata.cn/gene/) databases respectively, and then used as the query to search against the local wheat proteins by BLASP tool [53] with the identity more than 50% and e-value of 1e-10 as threshold. For BLASTP results, entries with the least expectation and the highest coverage were considered as the optimal target.

## Author’s contribution

N.X.J., C.L.C., Y.H, H.D.J. and D.P.C. designed and supervised the project. N.X.J., Y.G., P.Y. and C.L.C. collected and generated the data, and perform analysis. Z.Q.D. performed GWAS analysis. P.W.Q. and Z.L. contributed to data analysis. H.D.J. contributed to plant material collection and genotyping. N.X.J. and Y.G. prepared the draft manuscript. N.X.J., C.M.X, Y.H., R.J.H., D.E. and J.B. reviewed and revised the manuscript.

## Acknowledgments

This work was funded by Tang Scholar in NWSUAF, and also partially supported by the National Natural Science Foundation of China (Grant No. 31771778 and 31971885) and the Programmer of Introduction Talents of Innovative Discipline to Universities (Project 111) from the State Administration of Foreign Experts Affairs, China (Grant No. #B18042). We are grateful to High-Performance Computing center of Northwest A&F University for providing computational resources in this work.

## Data available

RNA sequencing data used in this study has been deposited into the Genome Sequence Archive (GSA) database with the accession number of PRJCA004969 and other data was provided in Supplemental information files.

## Supplemental information Supplemental Figures

Figure S1. Statistic of RNA-seq sequencing data and transcript assembly.

Figure S2. Genome-wide association studies of spike-related traits.

Figure S3. Co-expression network analysis identified the modules associated with spike development in wheat.

Figure S4. Genome-wide average LD decay estimated from 89 samples.

Figure S5. Characteristics of the identified sQTLs.

Figure S6. Co-expression network of hub genes in M3 module.

Figure S7. Co-expression network of hub genes in M5 module.

Figure S8. Co-expression network of hub genes in M4 module.

Figure S9. Co-expression network of hub genes in M7 module.

Figure S10. *TraesCS4B02G395700* and *TraseCS5A02G557300* were detected with GWAS signals that regulated spike length in SL-related M1 module.

Figure S11. The expression level of *TraesCS5D02G454700* was regulated by *AX-111930824* and the splicing variation of *TraesCS5A02G547700* was regulated by *AX-109874238* in M1 module.

Figure S12. Threshold of gene expression screening.

## Supplemental Tables

Table S1. GWAS signals associated with spike-related traits in wheat.

Table S2. Orthologues of the genes involved in wheat spike-related traits through GWAS analysis.

Table S3. GO, PO and TO enrichment analysis of the genes in different modules obtained from WGCNA analysis.

Table S4. Summary of sQTL signals.

Table S5. Summary of eQTL signals.

Table S6. Conserved protein domains variations in different isoforms of sGenes.

Table S7. Change of miRNA-binding sites in different isoforms of sGenes.

Table S8. Overlap between cis-sQTLs and cis-eQTLs.

Tables S9. Overlap between cis-eQTLs and GWAS signals.

Table S10. Overlap between cis-sQTLs and GWAS signals.

Table S11. Overlap between cis-sQTLs, cis-eQTLs and GWAS signals.

Table S12. KEGG enrichment analysis of different WGCNA modules.

Table S13. Key genes in the molecular module associated with grain number per plant.

Table S14. Key genes in the molecular module associated with spike length.

