## Supplemental Figures for "Genetic basis of expression and splicing underlying spike architecture in wheat (*Triticum aestivum* L.)"

**Supplementary Figures**

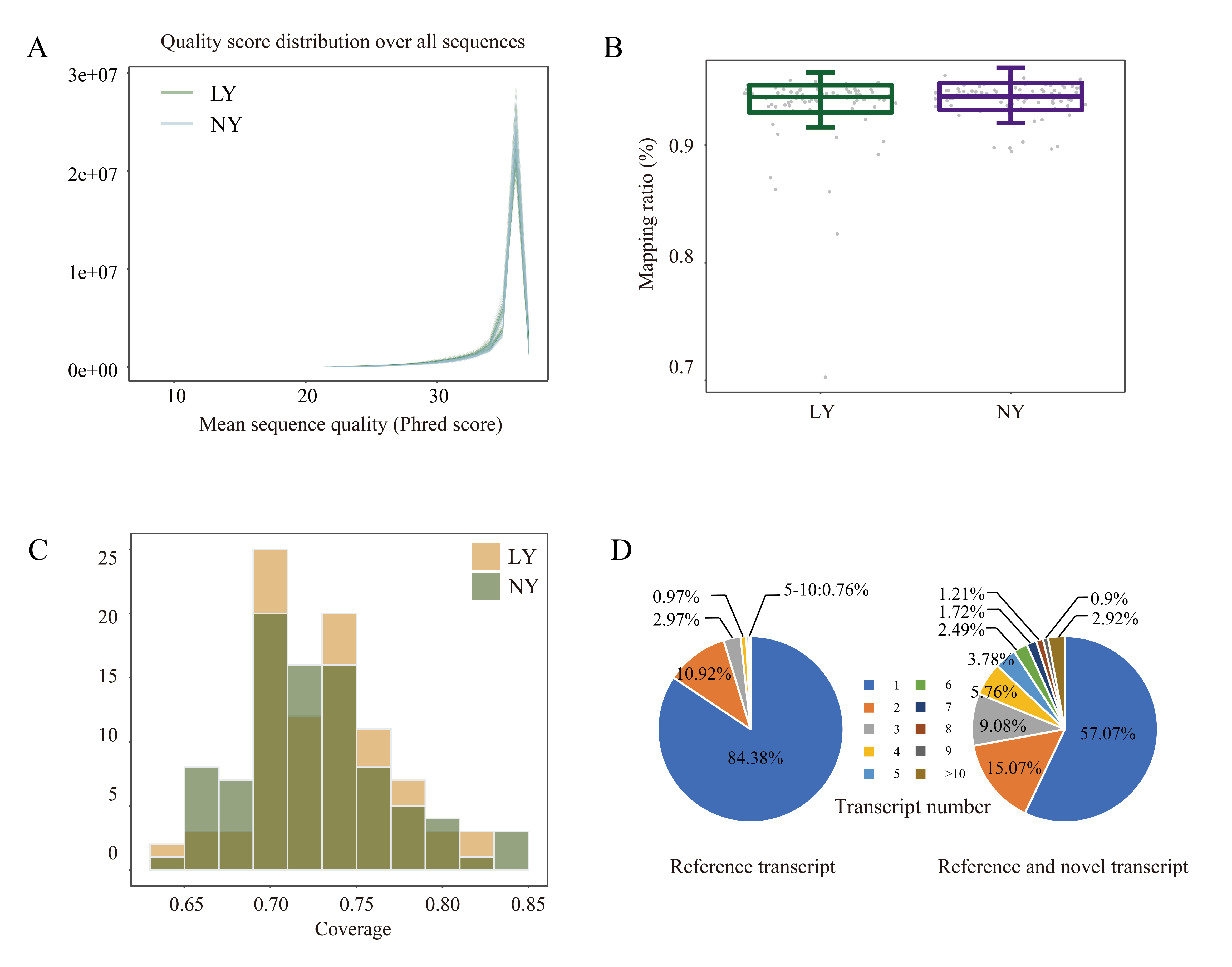

**Figure S1. Statistic of RNA-seq sequencing data and transcript assembly.**

**(A)** Sequencing reads quality of each sample in LY and NY shows the majority of our reads were of high quality.

**(B)** Mapping ratio of all samples in LY and NY shows the majority of our reads were mapped to reference genome.

**(C)** Relative sequencing reads coverage across the genome, showing good coverage along chromosomes.

**(D)** Distribution of transcript isoform number before and after assembling, showing the multi-isoform genes increased from 15.6% to 42.9%.

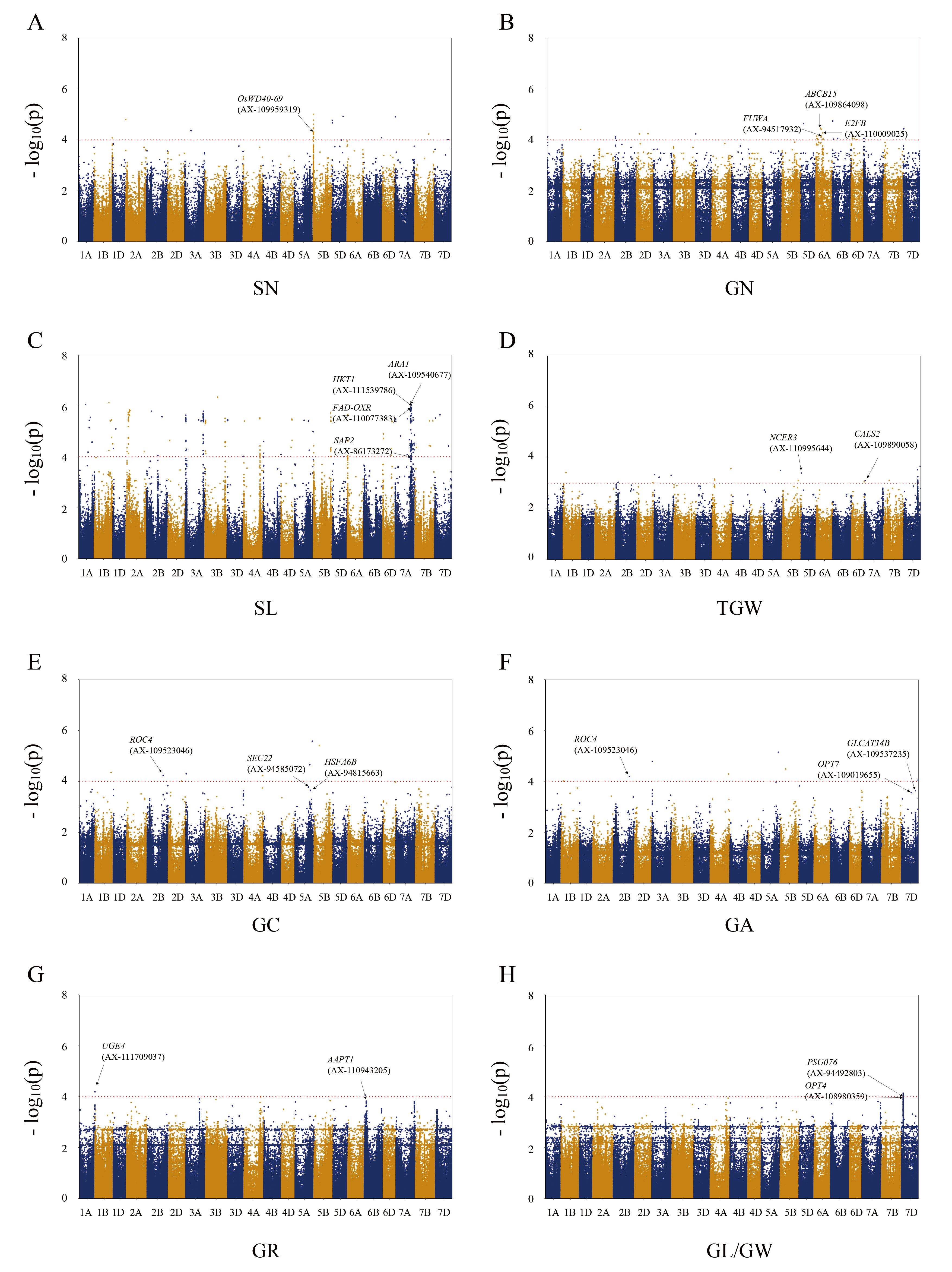

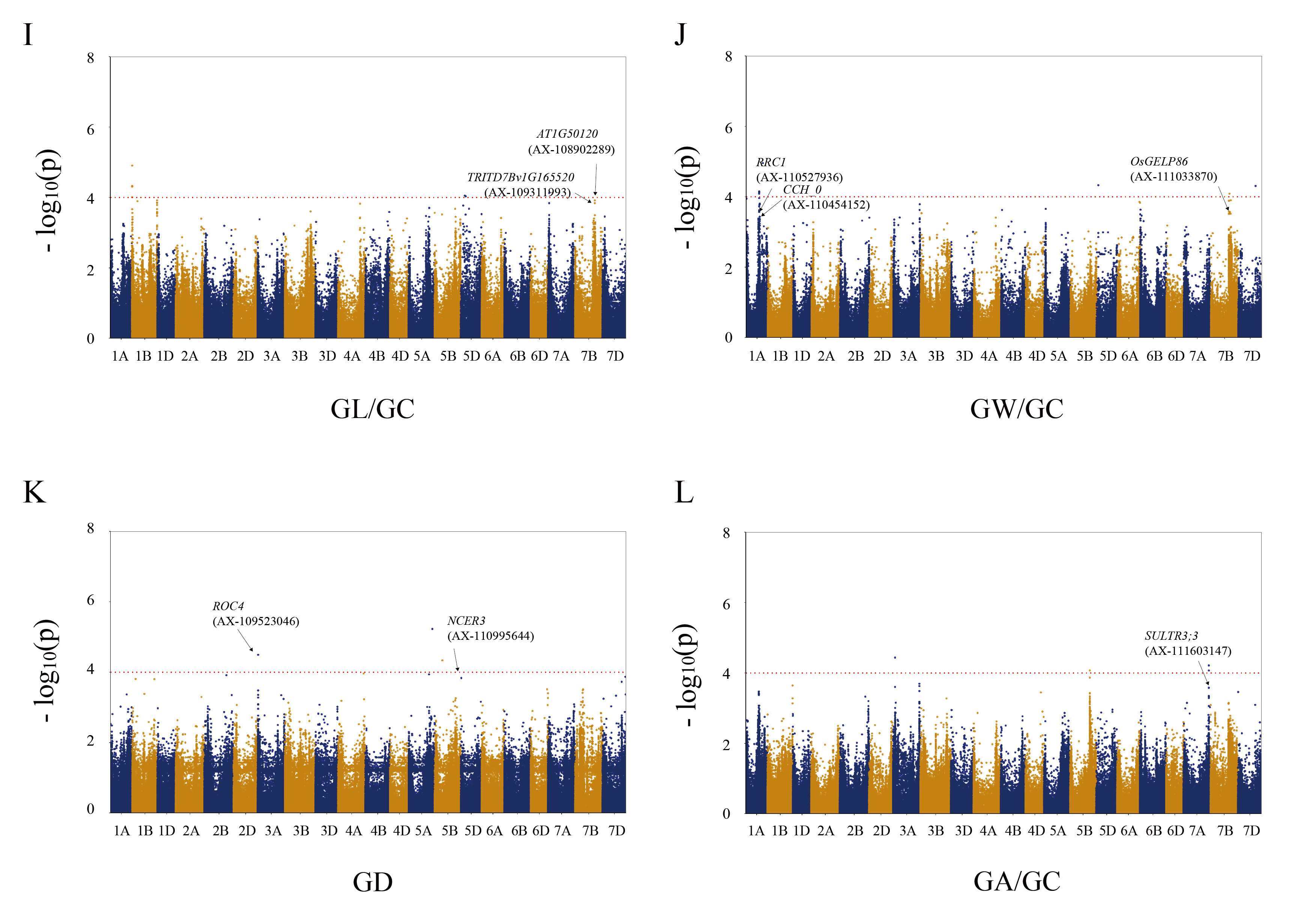

**Figure S2. Genome-wide association studies of spike-related traits.** Manhattan plots of compressed MLM for spikelet number (A), grain number (B), spike length (C), thousands grain weight (D), grain circumference (E), grain area (F), grain roundness (G), ratio of grain length to grain width (H), ratio of grain length to grain circumference (I), ratio of grain width to grain circumference (J), grain diameter (K) and ratio of grain area to grain circumference (L).

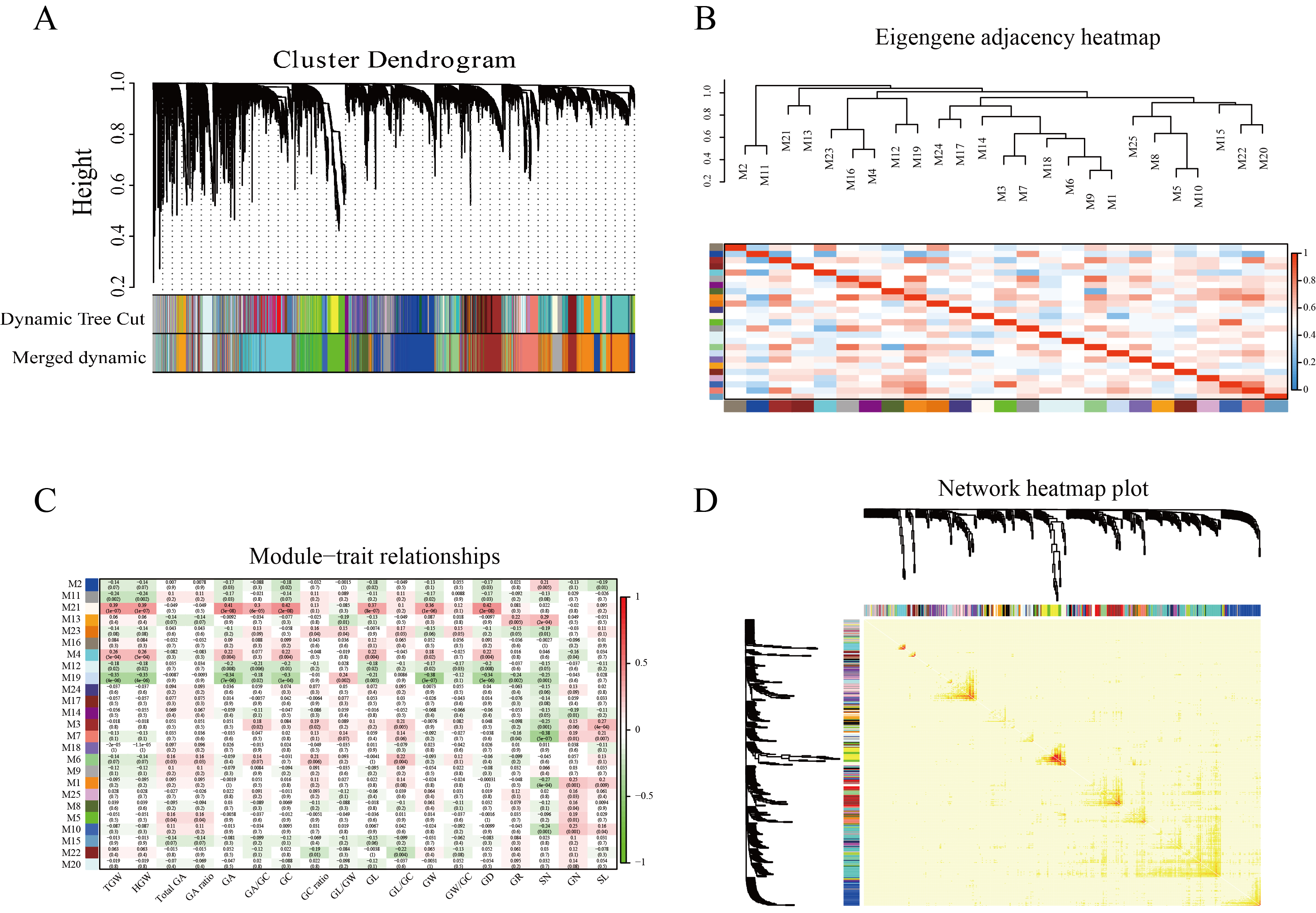

**Figure S3. Co-expression network analysis identified the modules associated with spike development in wheat.**

**(A)** Network analysis dendrogram based on hierarchical of genes by their topological overlap, identifying 25 co-expression modules. The color below shows the modules initially divided and the modules merged with high correlation. The modules are numbered M1-M25 in descending order of gene number.

**(B)** Heatmap of correlation between co-expression modules.

**(C)** Correlation between co-expression modules and traits. Negative value represents negative correlation, positive value represents positive correlation.

**(D)** A total of 1000 genes were randomly selected to construct network heatmap plot, high co-expression interconnectedness was indicated by progressively more saturated yellow and red colors.

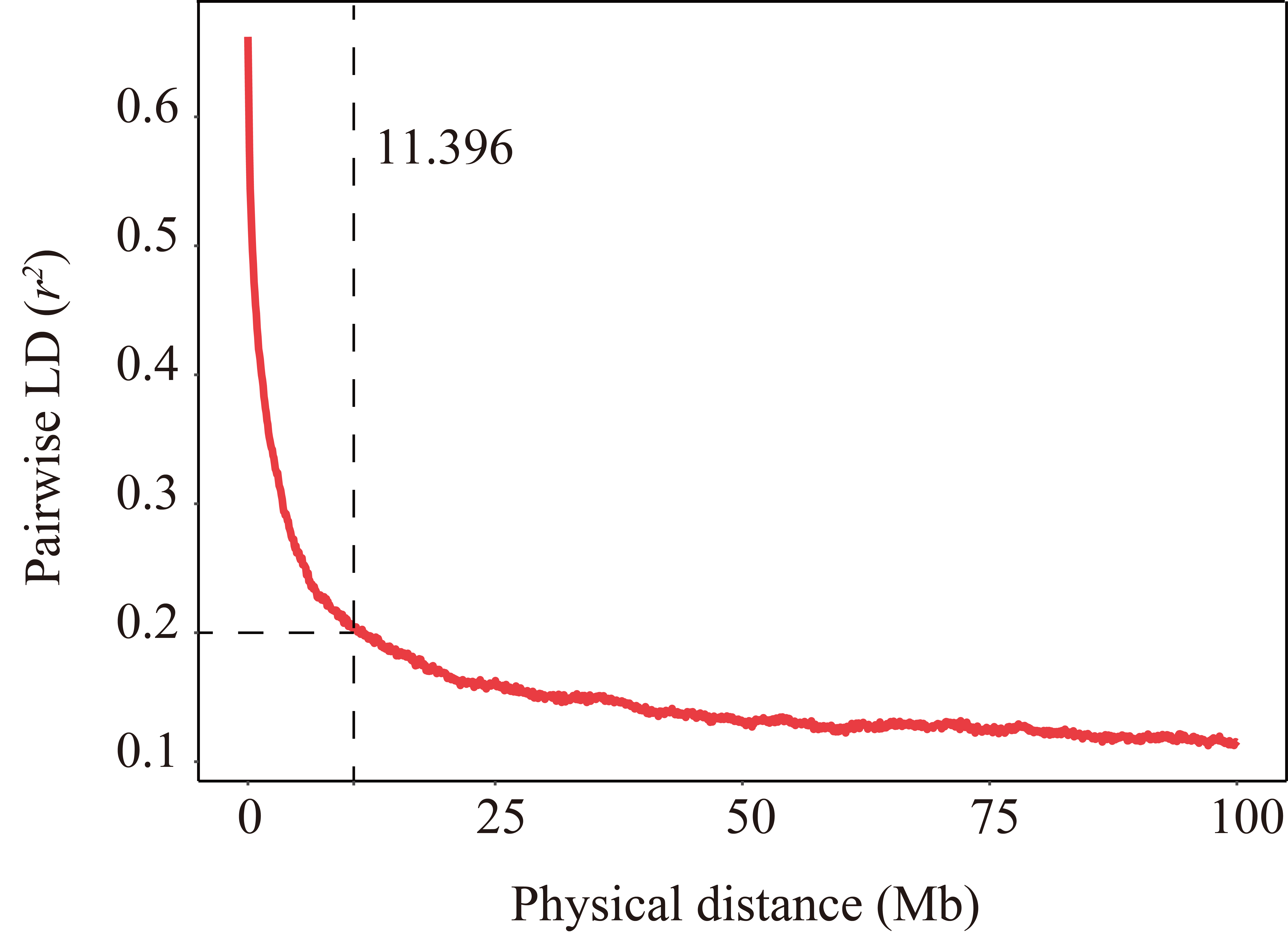

**Figure S4. Genome-wide average LD decay estimated from 89 samples.**

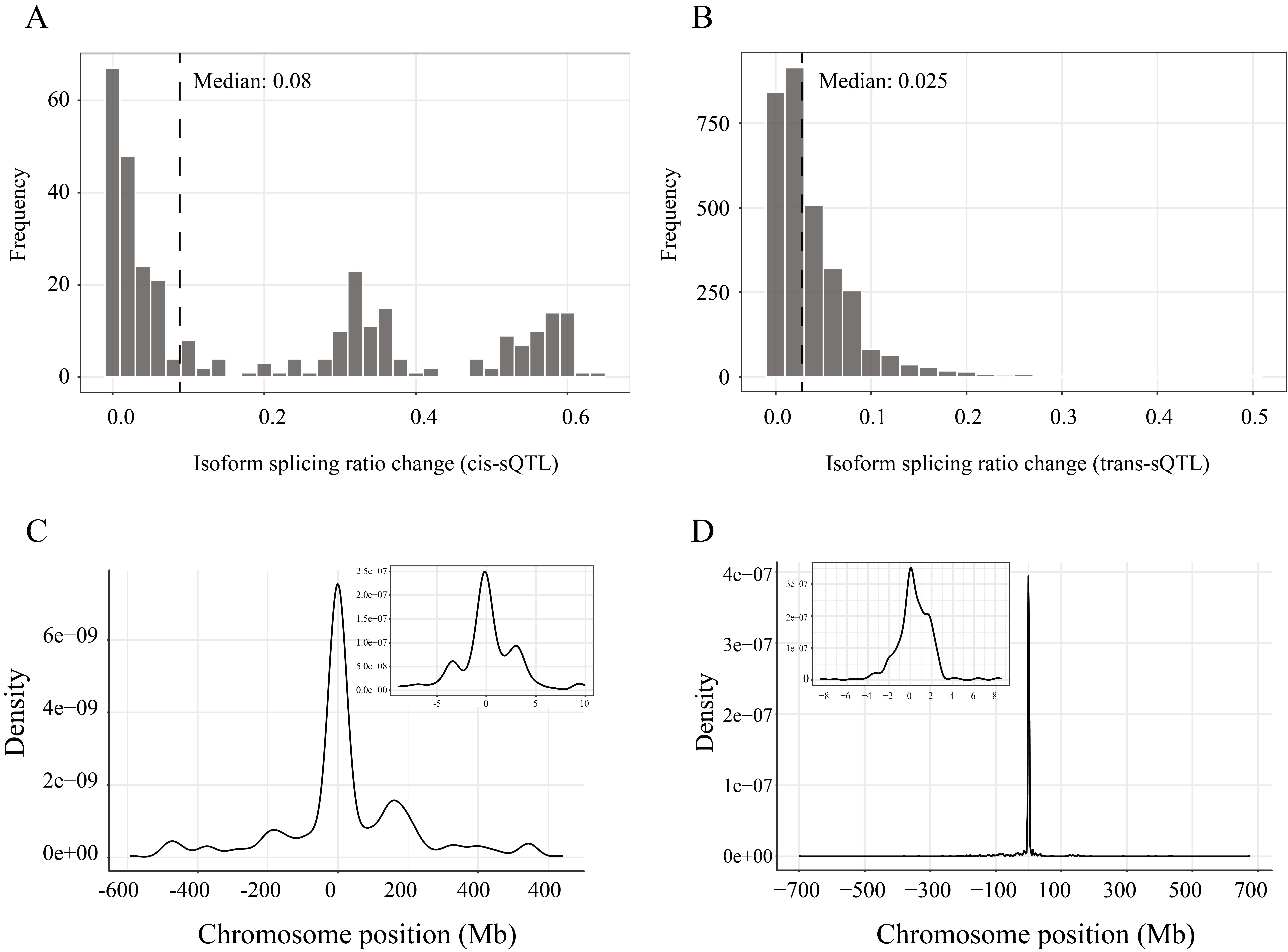

**Figure S5. Characteristics of the identified sQTLs.**

**(A)** Distribution of splicing ratio differences between genotypes at each cis-sQTLs.

**(B)** Distribution of splicing ratio differences between genotypes at each trans-sQTLs.

**(C)** Position enrichment of cis-sQTL based on TSS (Transcription start site).

**(D)** Position enrichment of cis-eQTL based on TSS (Transcription start site).

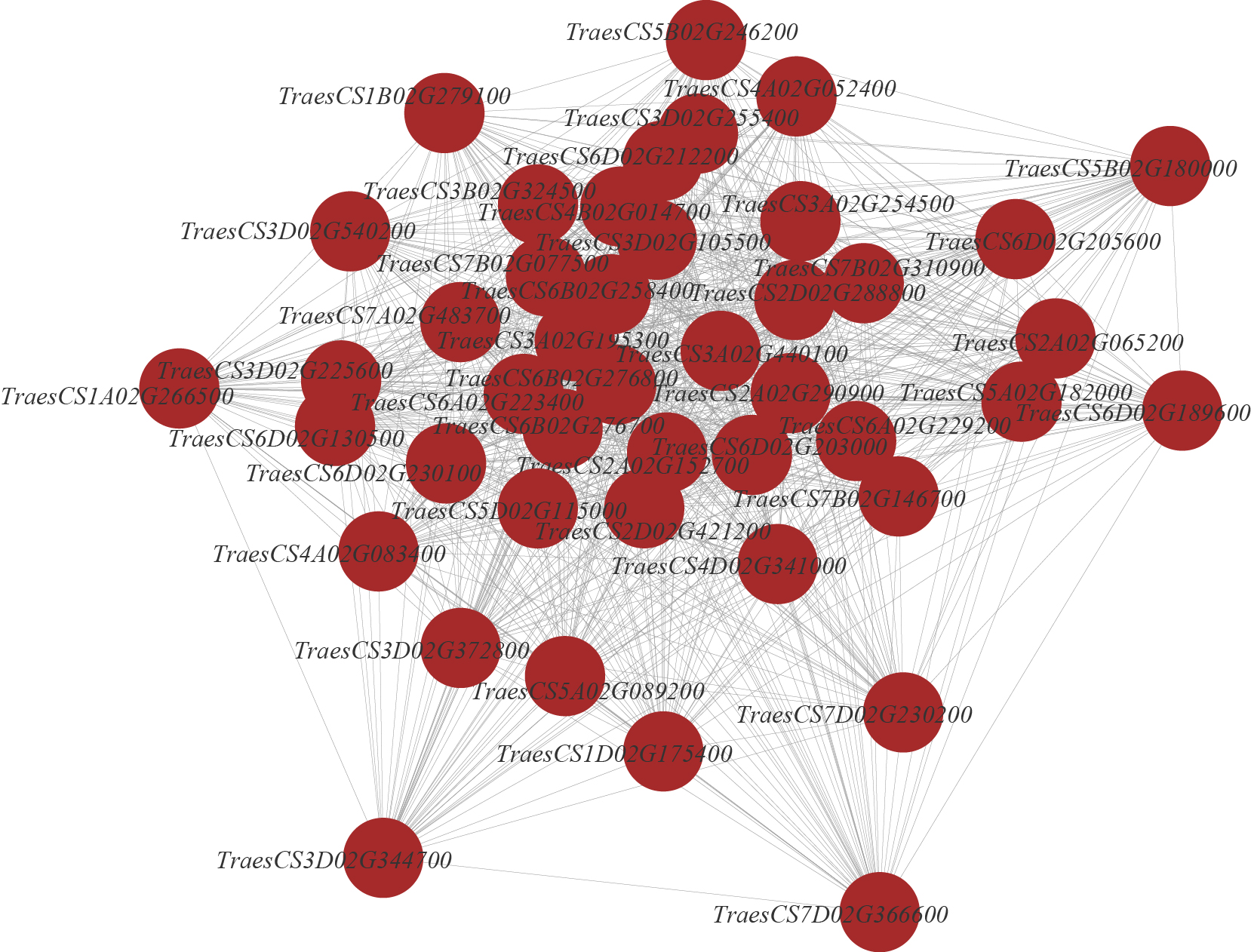

**Figure S6. Co-expression network of hub genes in M3 module.**

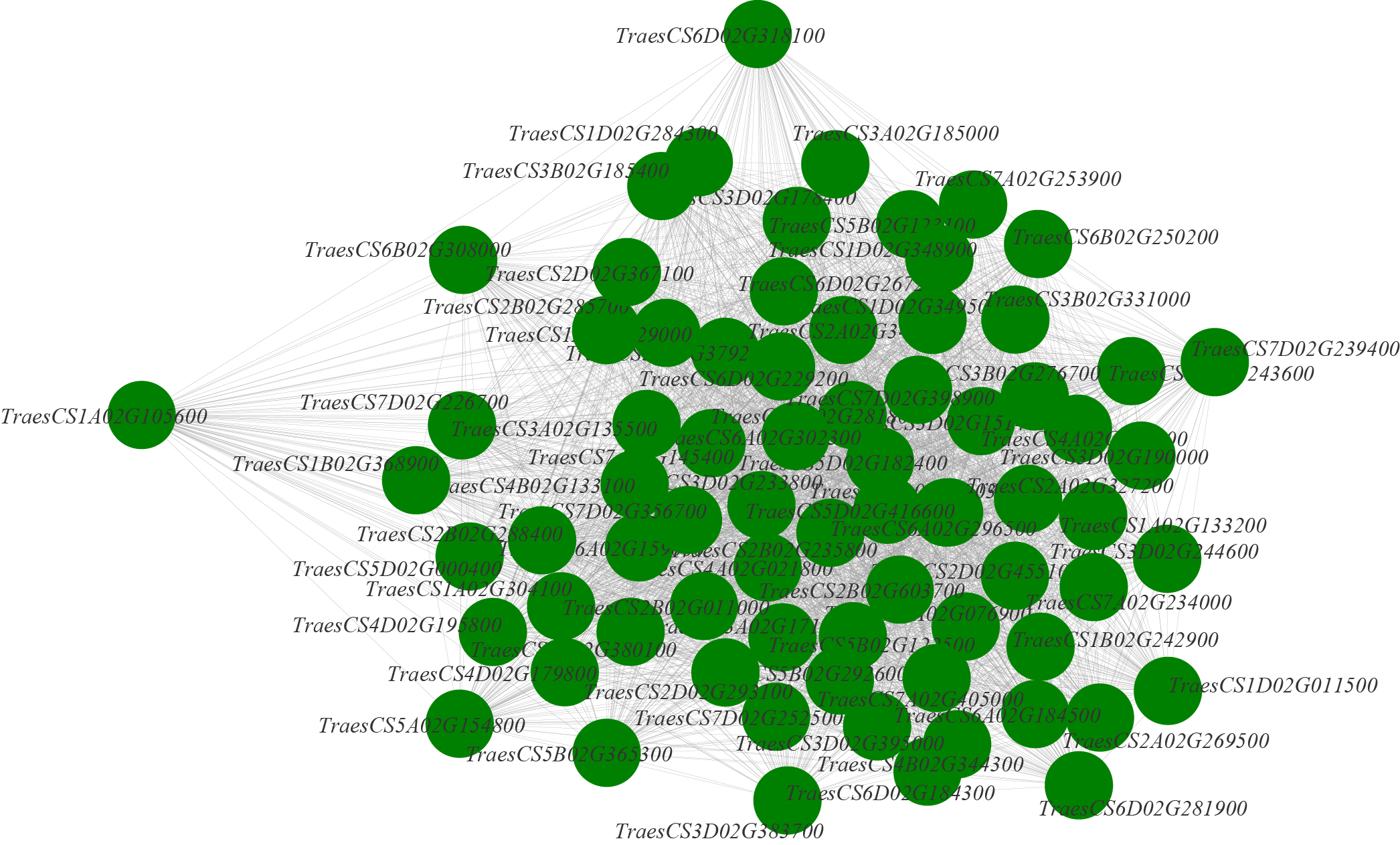

**Figure S7. Co-expression network of hub genes in M5 module.**

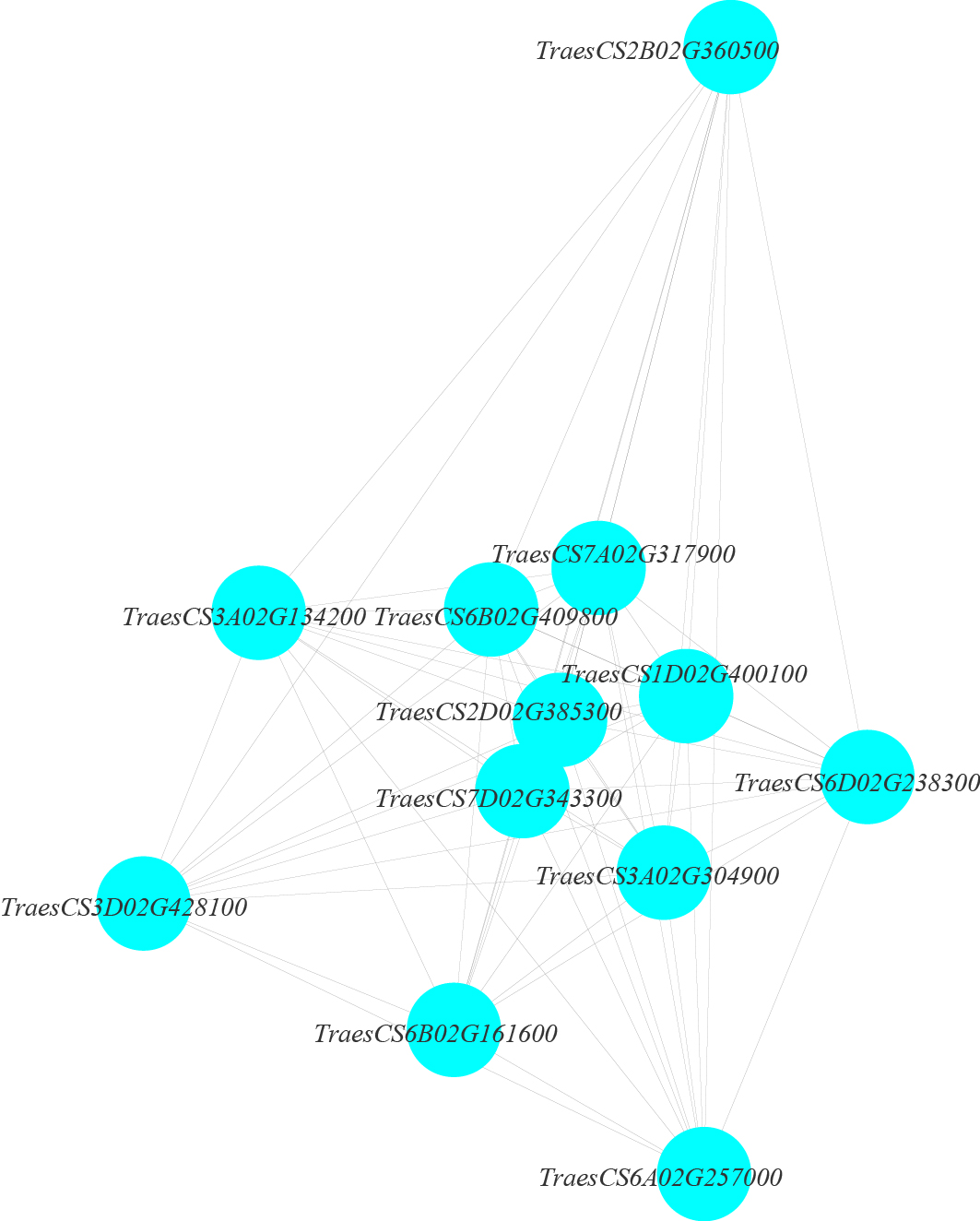

**Figure S8. Co-expression network of hub genes in M4 module.**

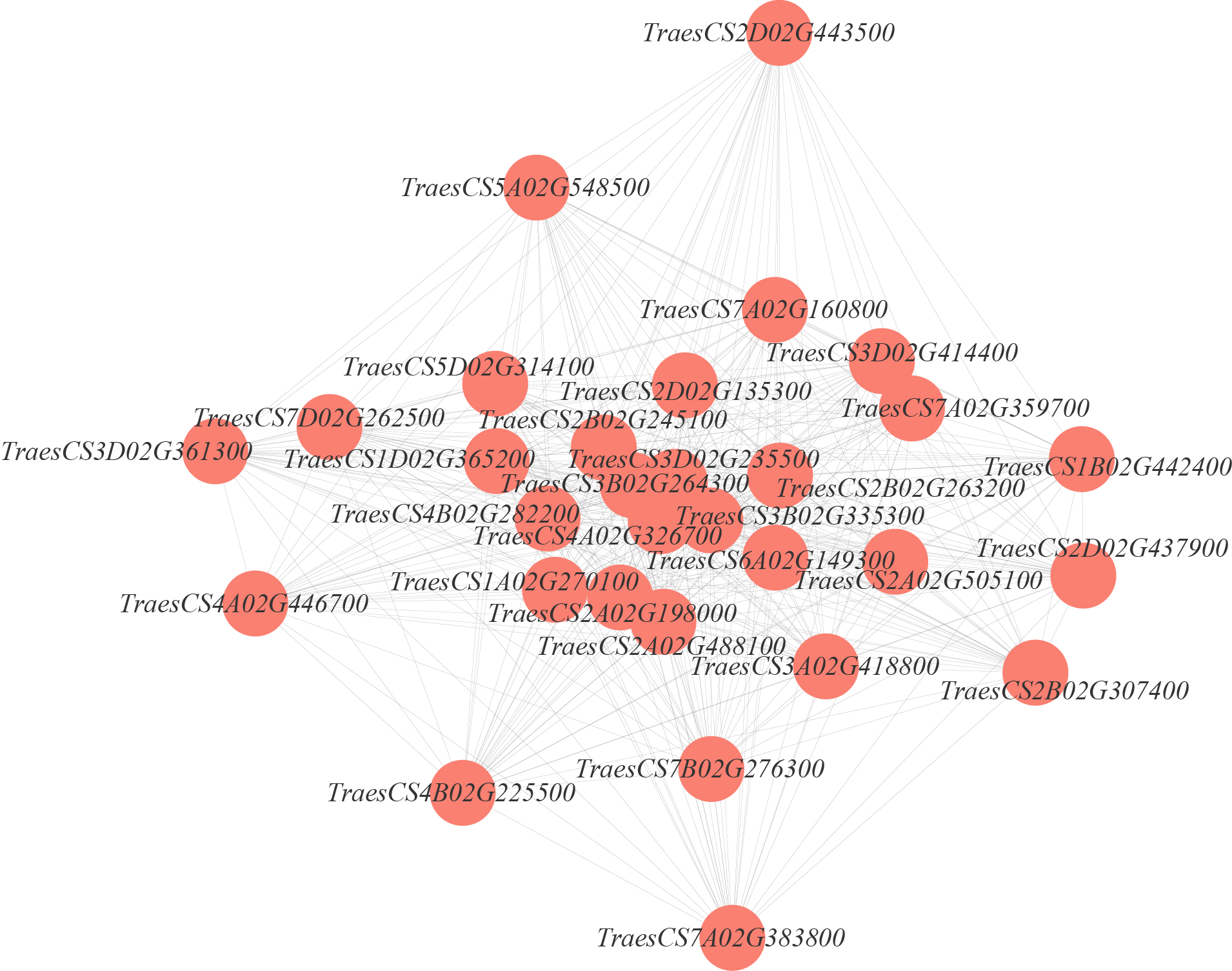

**Figure S9. Co-expression network of hub genes in M7 module.**

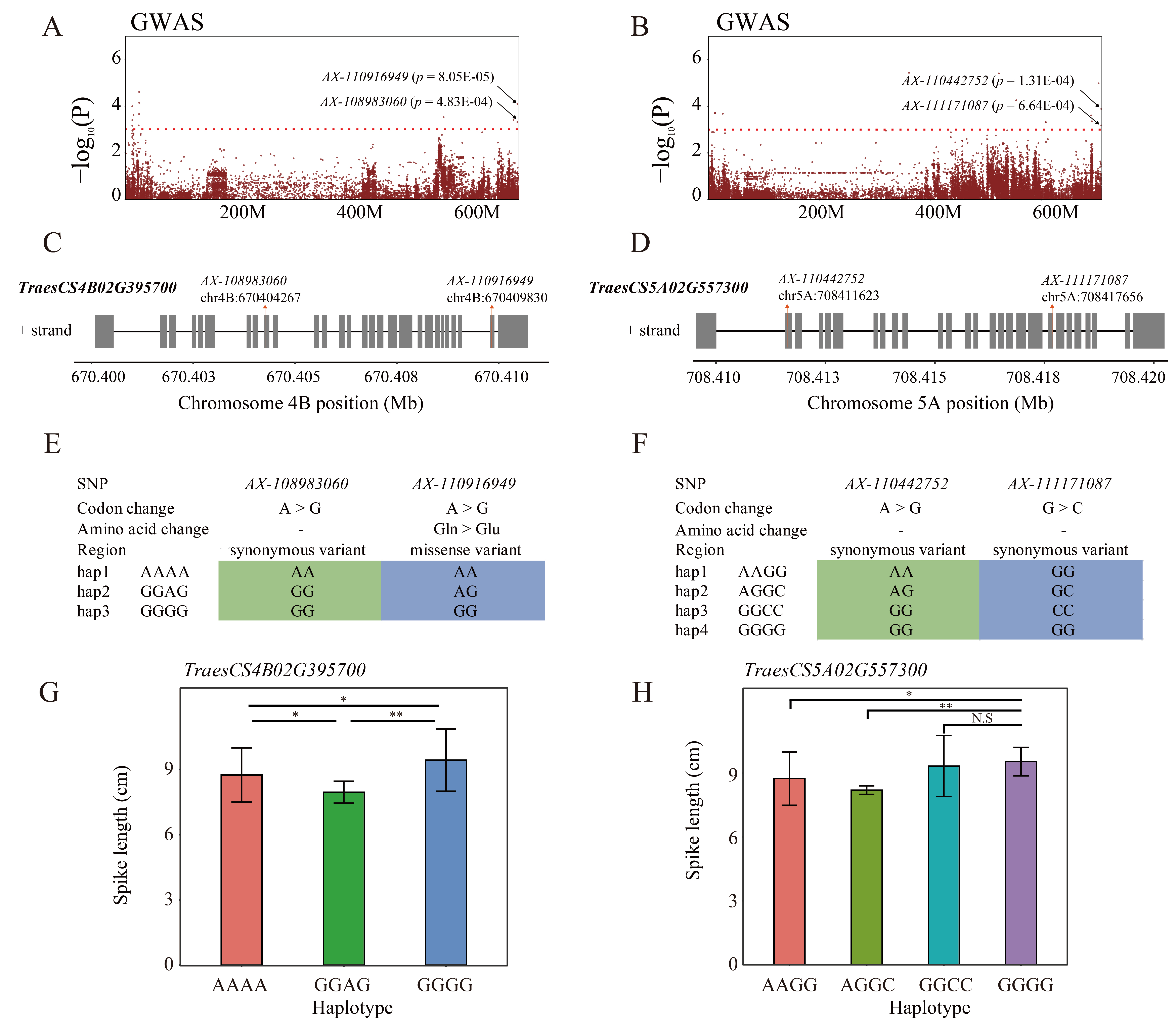

**Figure S10. *TraesCS4B02G395700* and *TraseCS5A02G557300* were detected with GWAS signals that regulated spike length in SL-related M1 module.**

**(A)** Manhattan plot of GWAS signal in *TraesCS4B02G395700*.

**(B)** Manhattan plot of GWAS signal in *TraseCS5A02G557300*.

**(C)** GWAS signals were located in exon-intron structures of *TraesCS4B02G395700.*

**(D)** GWAS signals were located in exon-intron structures of *TraseCS5A02G557300*.

**(E)** Haplotype in the gene region and the core haplotype were screened and analyzed in *TraesCS4B02G395700*.

**(F)** Haplotype in the gene region and the core haplotype were screened and analyzed in *TraseCS5A02G557300*.

**(G)** Effects of different haplotypes of *TraesCS4B02G395700* on spike length.

**(H)** Effects of different haplotypes of *TraseCS5A02G557300* on spike length. *, *p*< 0.05; **, *p*< 0.01; ***, *p*< 0.001; N.S, not significant.

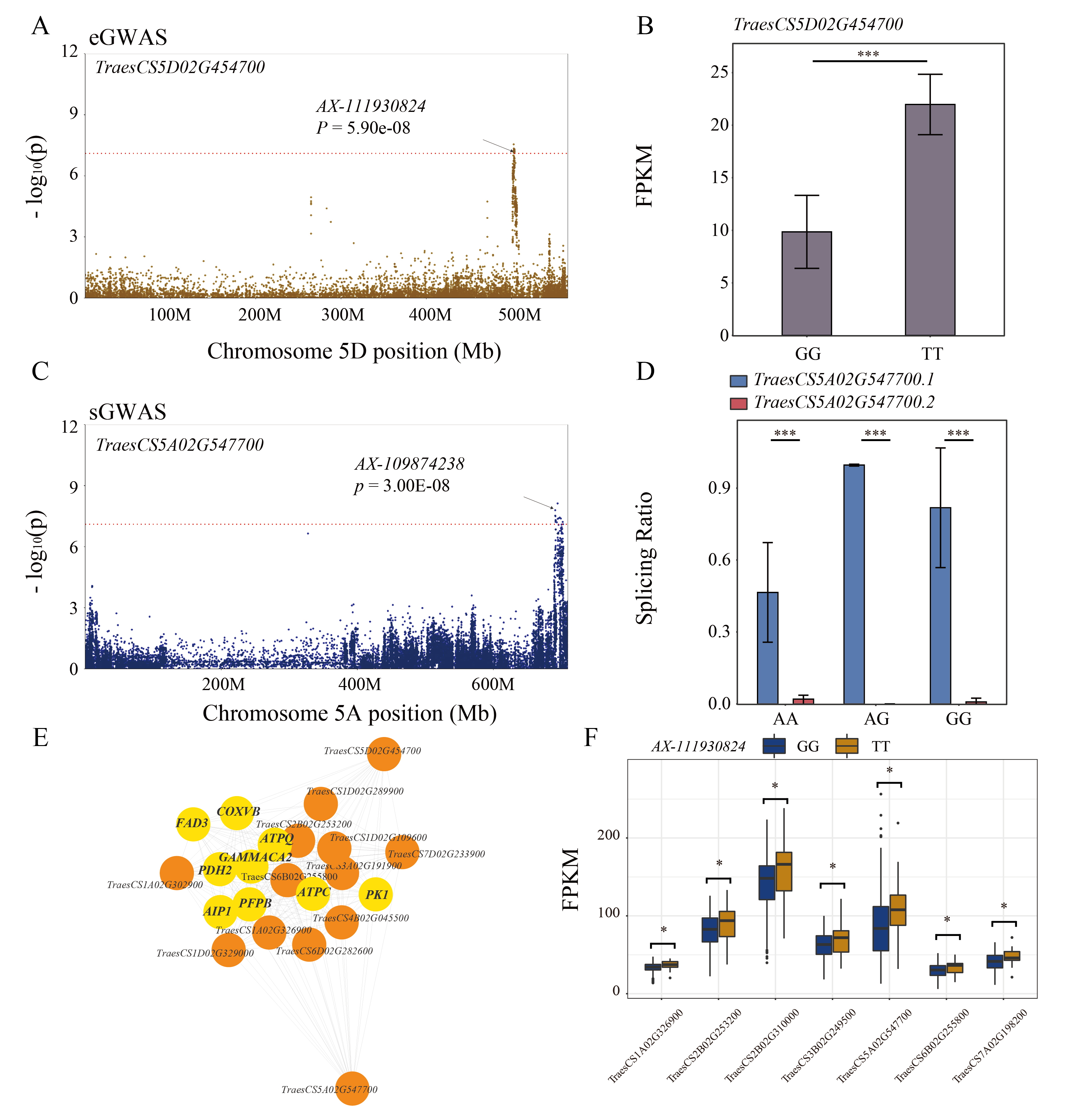

**Figure S11. The expression level of *TraesCS5D02G454700* was regulated by *AX-111930824* and the splicing variation of *TraesCS5A02G547700* was regulated by *AX-109874238* in M1 module.**

**(A)** Manhattan plot of eGWAS on 5D to show the relationship between the SNP loci with the expression level (FPKM value) of *TraesCS5D02G454700*.

**(B)** Gene expression variations among different genotypes.

**(C)** Manhattan plot of sGWAS 5A to show the relationship between the SNP loci with the alternative splicing ratio (AS Ratio) of *TraesCS5A02G547700.*

**(D)** Gene alternative splicing variations among different genotypes.

**(E)** Co-expression network in M1 module.

**(F)** Expression variations of co-expression genes highly linked to eGene and sGene in M1 module. *, *p*< 0.05; **, *p*< 0.01; ***, *p*< 0.001; N.S, not significant.

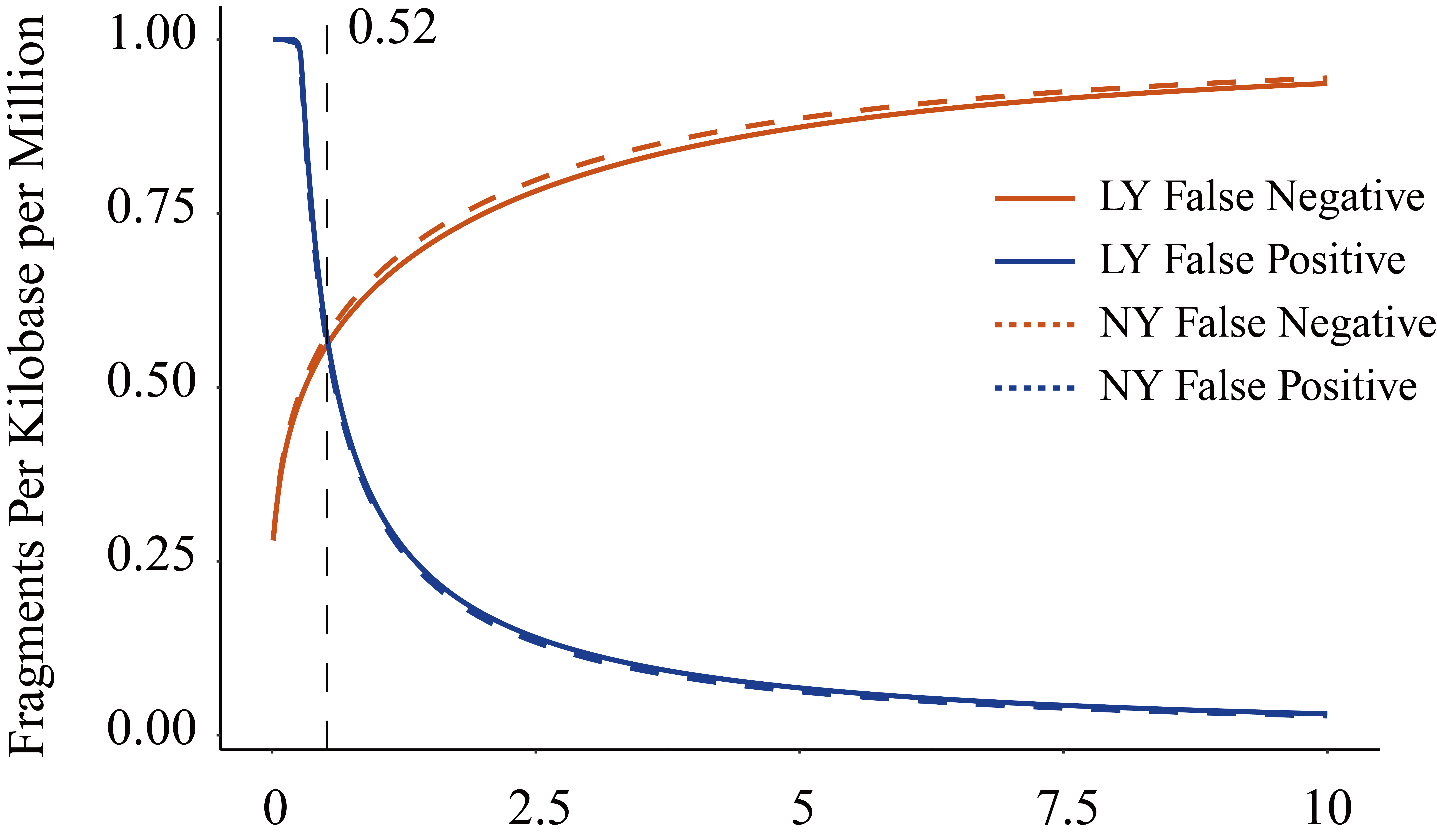

**Figure S12. Threshold of gene expression screening.** Distribution of reference transcripts with low or absent expression (False Negative) plotted against newly assembled transcripts with high expression (False Positive). The proportion of false negative was higher than that of false positive FPKM as cutoff, and isoforms needs to exceed the threshold in at least 5% of the samples.
